# Functional Group Composition: The Blueprint for Protein Interactions

**DOI:** 10.1101/2025.10.07.680707

**Authors:** Torbjörn Nur Olsson, Atsarina Larasati Anindya, María-José García-Bonete, Elwin Vrouwe, Didier Marguet, Jordan Preto, Vania Calandrini, Marco Pettini, Sandra Ruffenach, Jeremie Torres, Venkataragavan Chandrasekaran, Maria Bokarewa, Stefano A. Mezzasalma, Gergely Katona

## Abstract

Understanding the complex landscape of protein interactions, especially those involving intrinsically disordered proteins (IDPs), is fundamental yet challenging due to their structural heterogeneity and flexibility. Traditional sequence-based homology methods frequently fall short in characterizing IDP functions and interactions. Here, we present a novel approach leveraging supervised and unsupervised machine learning techniques, focusing exclusively on the compositional features of proteins. An Edmond-Ogston-inspired mixing model can reliably predict the degree of survivin (BIRC5) binding as a function of peptide composition alone, revealing a first interesting connection with the composition diagrams of chemical thermodynamics. By representing protein sequences through their functional group compositions, we demonstrate that specific compositions robustly predict binding interactions with survivin, an important human protein in cellular regulation pathways. Experimental validation via peptide microarray confirms the predictive power of our simplified compositional model, independent of exact amino acid sequences. Extending this method across the human proteome, we identified distinct compositional signatures correlating with survivin interactions and revealed fine grained biologically meaningful functional clusters based on compositional similarity. Our findings suggest a compositional blueprint underpinning protein interactions, offering a powerful, simplified framework to decode complex biological networks.

## Introduction

Protein functions span a spectrum from catalytic activities—driven by precise structural features—to broader biological processes such as cell division, subcellular localization, and programmed cell death. These functional domains often intersect, and proteins frequently display multifunctionality ^1^. This is particularly evident in cellular contexts such as mitosis or apoptosis, where many proteins assume multiple roles beyond strictly enzymatic activities like kinase or protease function. At a system level, cellular processes typically rely on coordinated actions from multiple proteins. Though, individual proteins often contribute to multiple, seemingly unrelated pathways—a phenomenon commonly referred to as moonlighting.

To link any kind of function to a protein sequence similarity metrics, sequence identity provides a good first approximation. Using sequence alignment for functional assignment assumes that proteins with similar sequences have similar structure and are likely to carry out similar functional roles within the cell ^2,3^. However, divergent evolution can blur the link between a sequence of proteins until a point that common ancestry is unrecognizable. It is also challenging to understand the necessity of sequence divergence, how these variations translate into altered function, and when they remain neutral to the protein function or the organism as a whole. Notably, even with substantial proteomic divergence, a mouse brain can seamlessly integrate human stem cells ^4^.

Protein function also depends on interactions with other proteins, especially for complex cellular processes, as protein roles are associated with the networks of protein-protein interactions (PPIs) they participate in. These interactions are essential for critical cellular functions, such as signal transmission, cell growth, differentiation, and programmed cell death, forming an intricate network known as the “interactome.”

When a protein has few close homologs or shows low similarity to known proteins, inferring its function becomes especially difficult using the sequence similarity approach ^5^. Proteins involved in the same higher-level process may share little sequence similarity and may not be homologous. These proteins may also consist of multiple regions (signal peptides, domains) with different degrees of independence in their functional allocation. These motifs and domains allow proteins to participate in different kinds of interactions (specific or fuzzy)^6-8^. Conversely, homologous proteins with similar sequences may fulfill distinct higher-level roles and be distributed in the interaction networks due to divergent evolution, making their common ancestry irrelevant for determining their higher-level function.

To assess the evolution of proteins it is good to develop a concept of what properties of proteins evolve and what is the feature landscape these properties define. Sequence is an obvious property and the current reviewed UniProt database ^9^ for the human proteome contains 20,363 genes, with a total sequence length of 11,277,205 amino acids for the corresponding proteins. This is also roughly equivalent to the total number of short reading frames across the entire proteome, specifically 15-amino-acid reading frames in our case ^10^. The small size of a single reading frame conceals a substantial variation as 15 amino acid long frame can represent 20^15^ = 3.3 × 10^19^ sequence variants when considering the 20 most common natural amino acids.

If we consider only the amino acid compositions rather than their sequence, the number of possible combinations drops dramatically. Instead of representing a specific order, we focus on the counts of each amino acid type within the sequence. The number of unique compositions obtained by distributing 20 amino acid types across 15 positions, based on a combinatorial argument, is 1.9 × 10^9^.

This number of compositions is still larger than the potential variation provided by the proteome. Comparing these two variant numbers indicates that coincidental compositional similarity is relatively likely and does not necessarily imply homology. Nonetheless, homology or sequence similarity may still underlie such compositional similarity, though composition alone is an unreliable and noisy measure for establishing homology. While the evolutionary history of a protein is valuable, it is tangential to the physical processes the protein participates in. While an evolutionary link often implies compositional resemblance—supporting the idea of shared function along lineages—it is difficult to separate this interpretation from other perspectives that associate homology with more complex, shared protein attributes (structure, surface complementarity, etc.).

Our focus on protein composition follows a productive scientific strategy that seeks to understand complex phenomena as compositions of interacting fundamental parts governed by basic rules. Composition is central across other scientific fields as well: the periodic table organizes elements by composition of subatomic particles, chemical formulas define compounds by their elemental composition, bulk properties such as miscibility also depend on chemical composition and the ratio of C/O atoms in organic molecules.

Regarding biology, the usefulness of compositional similarity has been demonstrated in proteins lacking sequence homology, which can often adopt similar structural folds ^11^. This suggests that specific amino acid compositions are predisposed to form certain stable configurations. Over evolutionary time, particular atomic arrangements consistently yield stable folds, resulting in recurring structural motifs across diverse proteins. Consequently, these common folds reflect a natural tendency toward stability, even in the absence of sequence similarity.

Many human proteins are named based on their composition, suggesting it is one of their most defining characteristics for example “arginine and glutamate-rich protein 1” (**ARGLU1**, Q9NWB6), “serine/arginine-rich splicing factor 6” (**SRSF6**, Q13247) or “acidic leucine-rich nuclear phosphoprotein 32 family member E” (**ANP32E**, Q9BTT0). Beyond folded proteins, about one-third of human proteins contain intrinsically disordered regions ^12^, characterized by compositional biases that allow substantial sequence flexibility without loss of function^13^. This compositional property enables diverse interactions, notably in transcription factors and DNA-binding proteins like linker histones ^14^. It was suggested that intrinsically disordered proteins can change their conformational ensemble properties, leading to diverse multivalent interactions ^15^. These interactions, which involve multiple binding sites between molecules, enable highly interconnected, network-like connections. This, in turn, affects liquid-liquid phase separation behaviors, a phenomenon that has also been shown to vary with the amino acid composition in independent studies ^16^.

The size of a system that is associated with a certain function can be rather small. Short signal peptides exhibit strong compositional biases, determining protein localization of the entire protein ^17-20^. Amino acid functional group composition alone effectively predicted survivin(BIRC5)-binding interactions^10,21^, suggesting that such binding may rely on simple compositional logic rather than sequence-specific properties. Applying this approach to the human proteome, we identified proteins with enriched peptides potentially binding survivin were included in a new peptide microarray to validate the earlier model.

Firstly, we test the composition-based survivin binding hypothesis in a new peptide microarray experiment using a set of human proteins with predicted binding properties and peptides with simple, systematically varied composition ^10^ (Figure 1). Secondly, we develop a thermodynamic model that quantitatively predicts survivin binding. Thirdly, we build a new hypothesis by moving away from the protein-centric view, systematically decomposing proteins into short segments, and exploring the connection between compositional signatures and functional descriptors. In this context, we aim to establish the necessary level of detail in the compositional profile of potential signaling segments for accurate binding prediction, functional association, or enrichment analysis. We examine functional insights gained from clusters containing protein segments that are exclusively predicted to bind survivin, as well as those containing segments that resemble survivin-derived sequences but are not necessarily predicted to bind survivin exclusively. Our findings based on a compositional blueprint that underpin protein interactions offer a powerful, simplified framework for decoding complex biological networks.

**Figure 1.**
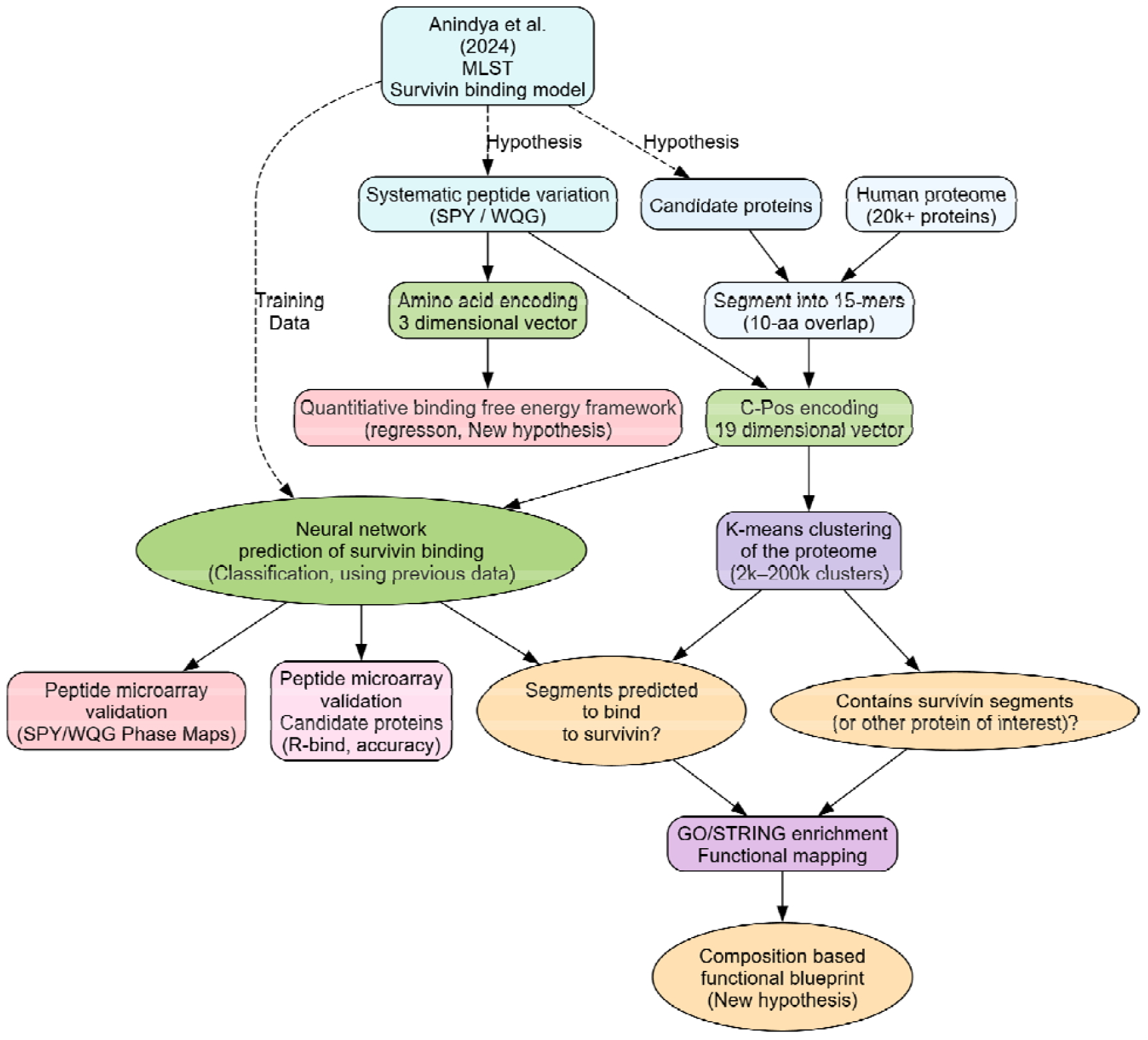
Overview of the study.

## Results and Discussion

### Validation of the composition-based model for survivin binding specificity

Experimental validation is necessary to assess whether computational models reflect underlying biochemical mechanisms. To test our machine learning model for detecting survivin-binding peptides ^10,21^, we performed a microarray experiment using peptides derived from human proteins predicted to be enriched in survivin-binding sequences, together with peptides of systematically varied composition.

As one set of experiments, we have chosen three amino acids (Serine-Proline-Tyrosine - SPY) and systematically varied these in blocks of five amino acids such that all possible composition (21 combinations) is present at least once in an arbitrary order of amino acids (Figures 2a). In another set of experiments, all possible permutations (total 243 variants) within five-amino-acid blocks were generated before replicating each sequence three times to produce 15-amino-acid peptides. The goal was to assess to what extent the amino acid order could be disregarded in favor of composition alone. For this analysis, we selected the amino acids tryptophan, glutamine, and glycine (WQG), as represented in Figure 2b. In Figure 2b, fluorescence values for permuted peptides with the same composition (21 variants) were averaged, before displaying as ternary diagrams.

**Figure 2.**
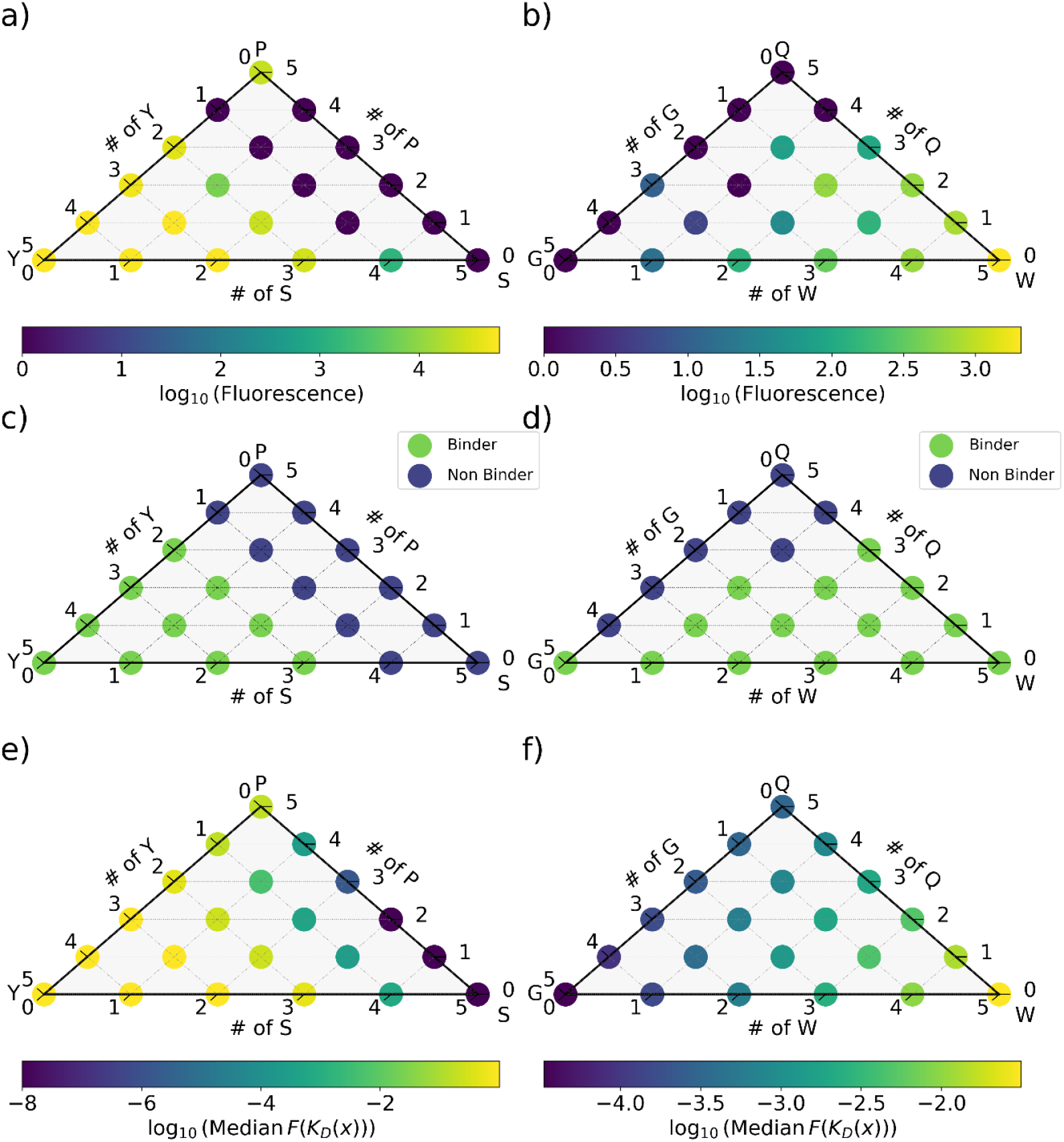
Ternary diagrams illustrating the effect of systematic variation in peptide amino acid composition on survivin binding. Panel a) and b) show experimental data from a survivin-binding microarray, represented as the logarithmic fluorescence intensity related to survivin binding (0 indicating no binding) for amino acid combinations of three amino acid residues serine, proline and tyrosine (S, P, Y) and tryptophan, glutamine and glycine (W, Q, G), respectively. Panels c) and d) depict neural network predictions, where the model, trained on the data set from Jensen et al ^21,46^ and C-Pos encoding of peptides, predicts survivin binding ternary diagrams for the same amino acid combinations SPY or WQG. Panels e) and f) depict the median predictions of the thermodynamic model for the Gibbs free function described here, where the data set is represented by amino acid identities.

The blocks of five amino acid segments were repeated three times to yield 15 amino acid long peptides. The reasons for this strategy were fourfold. First, sequence variations in five-amino-acid segments are manageable in a microarray experiment with a few thousand parallel experiments. Second, we identified a positional weight effect in peptide microarray experiments, and repeating the same segment multiple times helps to average out the influence of amino acid position along the sequence. Third, the sequence is less affected by inhomogeneities, and the local concentration of functional groups is more evenly distributed compared to cases where segments are not repeated. For example, the C-terminal region of the **SSSSSYYYYY** peptide may produce the same type of signal as **YYYYYYYYYY**, whereas **SYSYSYSYSY** may generate a distinct signal that locally homogeneous peptides cannot replicate. We will discuss later this aspect in the frame of chemical mixtures and the application of the Edmond–Ogston model (EOM)^22^. Fourth, the only relevant factors of 15 are 3 and 5, with variation across five positions providing a more detailed ternary diagram.

Figure 2a and 2b presents experimental data, while Figure 2c and 2d displays neural network predictions (hypotheses) based on C-Pos compositional encoding trained on previous data^10^. The experimental values closely correlate with the hypothesis, particularly in the SPY experiment where fluorescence intensity sharply declines when the number of tyrosine residues falls below two in the repeated sequence blocks or 40% of the peptide. However, contrary to the hypothesis, peptides containing only proline or a mixture of 20% tyrosine and 80% serine remain sufficient to provoke survivin binding.

The WQG hypothesis differed from the SPY hypothesis in that serine and proline were expected to be equally detrimental to survivin binding at this level of ternary diagram granularity, while glutamine was predicted to be more detrimental than glycine in the WQG experiment. The classification did not differentiate between varying binding strengths; zero fluorescence was considered non-binding, while non-zero values were classified as binding. Experimentally, WQG peptides generally exhibited weaker binding than SPY peptides, yet the asymmetry in the ternary diagram was confirmed, even if it did not perfectly align with the hypothesis. For instance, 80% glycine was better tolerated by survivin, whereas 80% glutamine was not when the remaining amino acid were tryptophane residues.

In the WQG experiment not all peptides with the same composition behaved identically at the individual level. One possible explanation, as previously mentioned, is the positional weight effect, an experimental artifact of the peptide microarray experiment ^10^. This effect can influence the amino acid at a given position and may even be noticeable between adjacent amino acids, particularly in the C-terminal region of the peptides. The neural network used is exceedingly simple, consisting of only two intermediate-layer neurons and 40 parameters, which limits the achievable topologies in the hypothetical ternary diagrams. However, this simplicity also reduces the likelihood of overfitting the noisy data. Moreover, local interaction patterns, charge, hydrophobicity and shape complementarity may further contribute to peptide selectivity which is not captured by the compositional description. It is important to note that none of the amino acids used in these two experiments (SPY and WQG) are charged, illustrating that the phase behaviour cannot be reduced to just simple electrostatic interactions between net charges. Finally, random experimental variation can lead to weakly reproducible binder and non-binder observations, particularly when fluorescence intensity is low.

Another source of discrepancies is the change in survivin concentration between the microarrays used for training and verification. In the training data, binding was observed at 1 µg/ml, whereas in the verification experiments, a concentration of 10 µg/ml was required to achieve qualitatively similar results. This difference in protein concentration may account for some of the observed inconsistencies especially for weaker interactions such as peptides containing W instead of Y. In summary, the topology of experimental ternary diagrams aligns with expectations derived from a simple composition-based model, supporting the hypothesis wherein the amino acid order can be ignored as first approximation.

Naturally occurring protein segments often contain more than three types of amino acids, with their diversity typically referred to as “high complexity”. To evaluate peptides that resemble natural sequences more closely, we generated hypotheses based on all proteins in the human proteome, assessing their fraction of predicted survivin-binding segments ^10^.

For this analysis, we examined 15-amino-acid segments of each protein with a 10-amino-acid overlap, mirroring the approach used in a peptide microarray experiment. Classification was then performed using the trained model. We identified proteins with a high fraction of survivin-binding segments (using the quantity R_bind_ in ^10^) and included additional proteins in the verification set. The hypothetical R_bind_ and experimentally verified R_bind_ values are presented in Table 1, along with protein-specific accuracy values based on the classification accuracy of individual peptides within each protein.

**Table 1.**
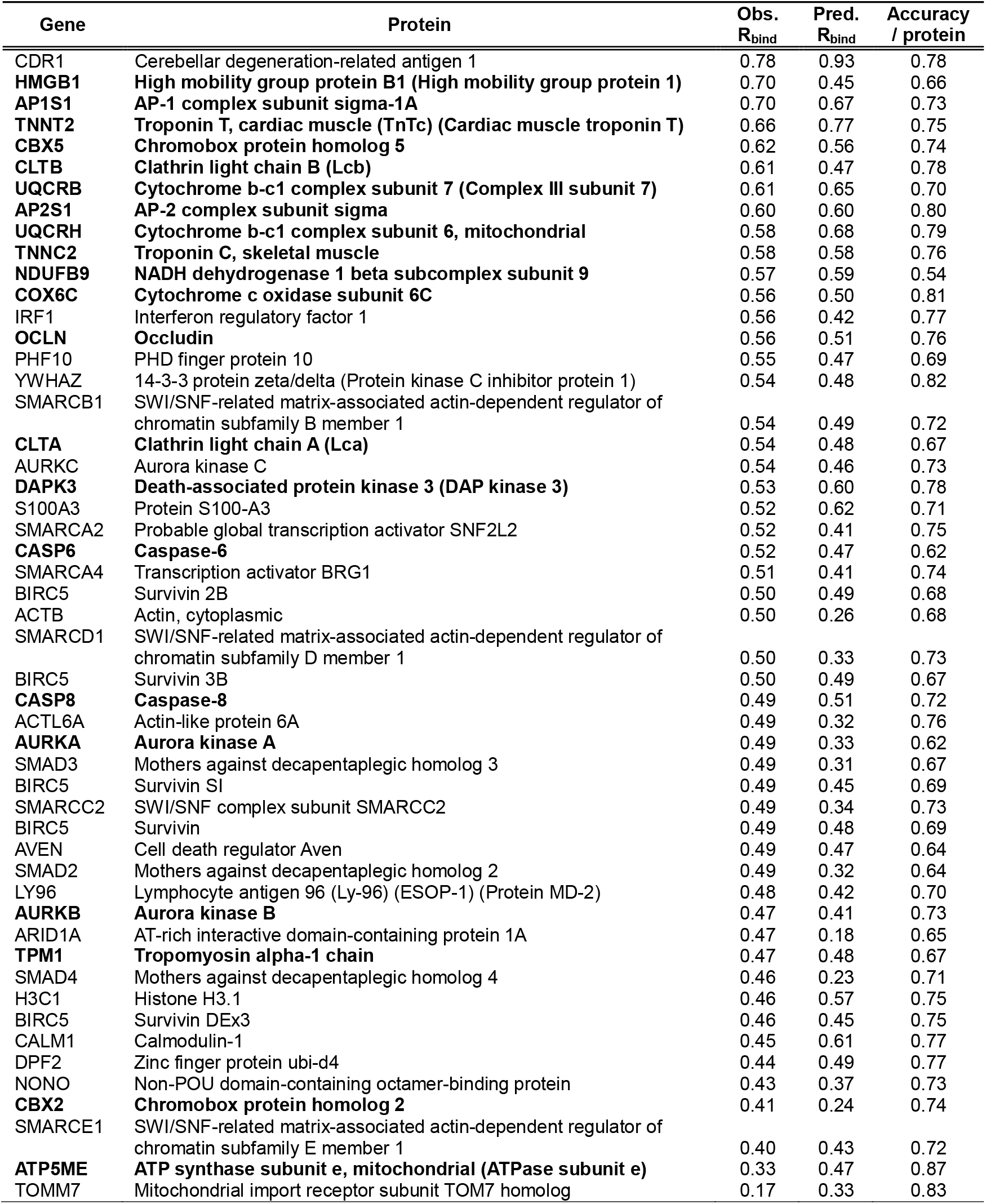
Fraction of survivin binding peptides (R_bind_) for selected human proteins and its prediction. The entries are sorted based on the observed R_bind_ values. Proteins in bold were explicitly mentioned in ref. ^10^.

The observed R_bind_ values exhibit a more compressed range than the predicted ones. This may result from extreme high and low predicted R_bind_ values having a higher proportion of false binder and false non-binder predictions than proteins with more typical values, respectively. Consequently, the actual values tend to shrink toward more typical levels. However, the ranking of proteins remains largely unchanged. Cerebellar degeneration-related antigen 1 retained the highest R_bind_ among the examined proteins, with an observed value of 0.78 even though this value is substantially lower than the predicted 0.93. Conversely, CBX2 was predicted to have only 0.24 of its peptide segments binding to survivin, but observed 0.41 during verification, still positioning it at the lower end of the observed R_bind_ scale. It is important to note that CBX5 was correctly predicted to have a much higher R_bind_ than CBX2 ^10^. The Pearson correlation between predicted and observed R_bind_ values in Table 1 was 0.60.

Summarizing this part, we have verified that peptides bind to survivin based on simple and generic composition-based models within a limited set of peptides. The predicted topology of these interactions is not highly complex, suggesting that survivin may recognize a broad range of compositions.

### Quantitative predictions from a thermodynamics-inspired Bayesian framework

Figure 2 presents ternary maps that inspired the development of a new quantitative model guided by thermodynamic principles. This equilibrium model associates the binding free energy change (ΔG_bind_) with the proportions of amino acids in the peptide partner. For simplicity, amino acid identities were used directly, as the three amino acids differ markedly in composition; however, the same principle applies when using chemical group representations in the feature vector.

The Edmond-Ogston ^22^ inspired model includes self-interaction, cross-interaction, and entropic contributions to ΔG_bind_. This free energy change determines the partitioning between bound and unbound forms of survivin, given protein and peptide concentrations, via the van’t Hoff relationship between ΔG_bind_ and the dissociation constant (K_D_).

Posterior parameters were inferred using a hierarchical Bayesian model, assuming normally distributed data with a half-normal prior on the scale parameter and uniform priors on three *B*_*ii*_ self⍰virial and three *B*_*kh*_ cross-virial coefficients. As shown in Figures 2e and 2f, the median of the posterior predictions qualitatively aligns with the observed ternary diagrams. Figure S1 further confirms a high quantitative correlation, with coefficients of 0.974 and 0.968 for the SPY and WQG datasets, respectively.

Such high correlations may reflect underlying simple patterns in the data but also suggest the possibility of overfitting. Indeed, as illustrated in Figures S2 and S3, several parameters exhibit broad posterior distributions, which can vary substantially under slight changes to prior assumptions—yielding nearly equally plausible predictions. This highlights the ambiguity associated with an underdetermined model, the potential for simplification or the need for more data to facilitate model selection. This interpretation is supported by Figures S4 and S5, which show correlations between parameters in the pair plots of the posterior traces.

While this model allows considerable flexibility in fitting the data, it may also be inappropriate due to the introduction of unmotivated constraints. In particular, the sigmoid shape imposed by the law of mass action—and more generally by the equilibrium chemical model—enforces a specific functional form. This is less problematic when the partitioning spans a wide dynamic range, from oversaturation to lack of binding or repulsion. However, for the WQG dataset, which contains subtle differences, the assumed shape of the binding curve becomes critically important.

We also have to evaluate any potential mechanistic link to original EOM model itself. Each point in the ternary diagrams (Figure 2) represents an interaction state between a peptide of given composition (**x**) and the survivin macromolecule. In the EOM model, components translationally and rotationally diffuse freely and become concentrated or depleted in different spatial domains of a liquid mixture according to their predicted potential. In contrast, the present peptide system constrains components (functional groups or amino acid residues) to a fixed distribution at very high concentration due to the remarkable stability of peptide bonds keeping them together, even when their close association is entropically or enthalpically unfavorable. Another way to reconcile the physical system with the EOM model is to view the peptide as a chemical mixture containing both *mixed* (for example, **SYSYSYSYSY**) and *pure* segments (**YYYYYYYYYY**), or as an **S** chemical group *dispersed* in a pool of **Y** groups (**YYYYSYYSYY**). The nomenclature thus shifts from describing such sequences as *tandem repeats* or *low-/high-complexity* regions to the realm of chemical thermodynamics, where the distribution of components is more relevant than the exact order or sequence pattern of amino acid residues. Although amino acid residues cannot change their sequence positions to achieve more favorable (de-)mixing, peptides themselves and freely diffusible proteins such as survivin in our experiment can associate or (re-)fold according to their virial coefficients. The composition of survivin is not explicitly part of our model, but since it remains constant across all experiments, we focus on variations in the peptides. Nevertheless, a complete model in the spirit of the EOM framework should account for both interaction partners in a traditionally defined binary protein–protein interaction, while also incorporating the influence of the surrounding bulk solvent.

It can also be assumed that each fluorescence measurement reflects on the average of a conformational equilibrium. In other words, the virial coefficients in Eq. (4) can be equivalently interpreted as linked to the self-interactions between residues (i-i) and to the cross-interactions between (k-h), mediated by portions of survivin chain that are freely diffusing in solution, and whose conformational displacements contribute to changing the free energy of the local system. This is also consistent with the standard formulation, where all B_*ij*_ account for the interactions with the liquid medium. The only, yet essential, difference lies in the nature of the interacting chemical species. In the original EOM they are free, whereas here they are mediated by a free macromolecular system.

Another point worth noting is the system incompressibility constraint, expressed by the unitary conservation of amino acid fractions in Eq. (4). While this assumption, implying no free volume, is unnecessary in EOM, where the solvent is integrated out ^23^, here it provides greater clarity in parametrizing the ternary axis, since exact absolute concentrations in the microarray experiment are poorly defined for a number of reasons (site density, hydration, etc.). Conversely, precisely setting a reference energy value would be not only a tough task, but also inconvenient in our exploratory analysis, since it ultimately concerns comparisons between interaction states (virial coefficients). We have therefore arbitrarily adopted the 1 M standard state (V_0_ = 1660 Å^3^ per molecule), while reserving the option to examine the model reliability as a function of V_0_ in future studies.

Moreover, assuming equilibrium may be unrealistic in this context, as weakly bound survivin is likely removed during the washing steps of the microarray experiment. That said, this does not invalidate the qualitative insight provided by Eq.(4), where the self- and cross-interacting terms scale with the concentration or number of chemical groups. This new hypothesis is intended as a proof of concept and a suggested direction, rather than a definitive claim about the functional form of the thermodynamic equation. Additional data and variation of experimental parameters will be necessary to discriminate between models based on subtle differences in their performance.

### Survivin favors specific compositional signatures in the human proteome

We now shift our focus to the composition space represented by the human proteome. This space is approximately 1,000 times smaller than the theoretical maximum for 15-mer sequence segments. Notably, many of the compositions shown in Figure 2 never appear in the human proteome. This raises a key question: how different can peptides be within the human proteome while still being predicted to bind survivin? Conversely, how robust are the signals that do not attract survivin? Selectivity is crucial for key hub proteins such as survivin: it must activate pathways that promote cell proliferation while avoiding inadvertent activation of those leading to cell death. Consequently, activating interactions with pro-apoptotic proteins may need to be actively suppressed.

This question is crucial for two main reasons. First, living organisms experience non-detrimental mutations that also affect their composition, meaning that molecular organization mechanisms must be inherently robust to small compositional variation. Second, different proteins specialize in distinct functions, requiring specific structures and catalytic properties that depend on particular polypeptide sequences which in turn affect their composition. If sufficient compositional similarity between these functionally distinct proteins is not possible without compromising their roles, then such overarching compositional signals may not be suitable as organizational principles in practice.

The necessary granularity for functional specificity was assessed using K-means clustering of 15-amino-acid-long peptides derived from the human proteome, represented by their C-Pos encoding. A total of 2,214,715 protein segments were generated as previously described ^10^. Clustering was performed at three levels (k=2,000, 20,000, and 200,000 clusters) to analyze different degrees of granularity. Smaller clusters were expected to contain a more homogeneous set of groups. Then, we examined the fraction of peptides predicted to bind survivin within each cluster. The distribution of clusters based on the fraction of survivin-binding peptides is shown in Figure 3. Even in the coarse-grained representation (2,000 clusters), 50% of the clusters can be classified as clear survivin binders (0.9–1.0: 10%) or clear non-binders (0.0–0.1: 40%) based on survivin binding predictions.

**Figure 3.**
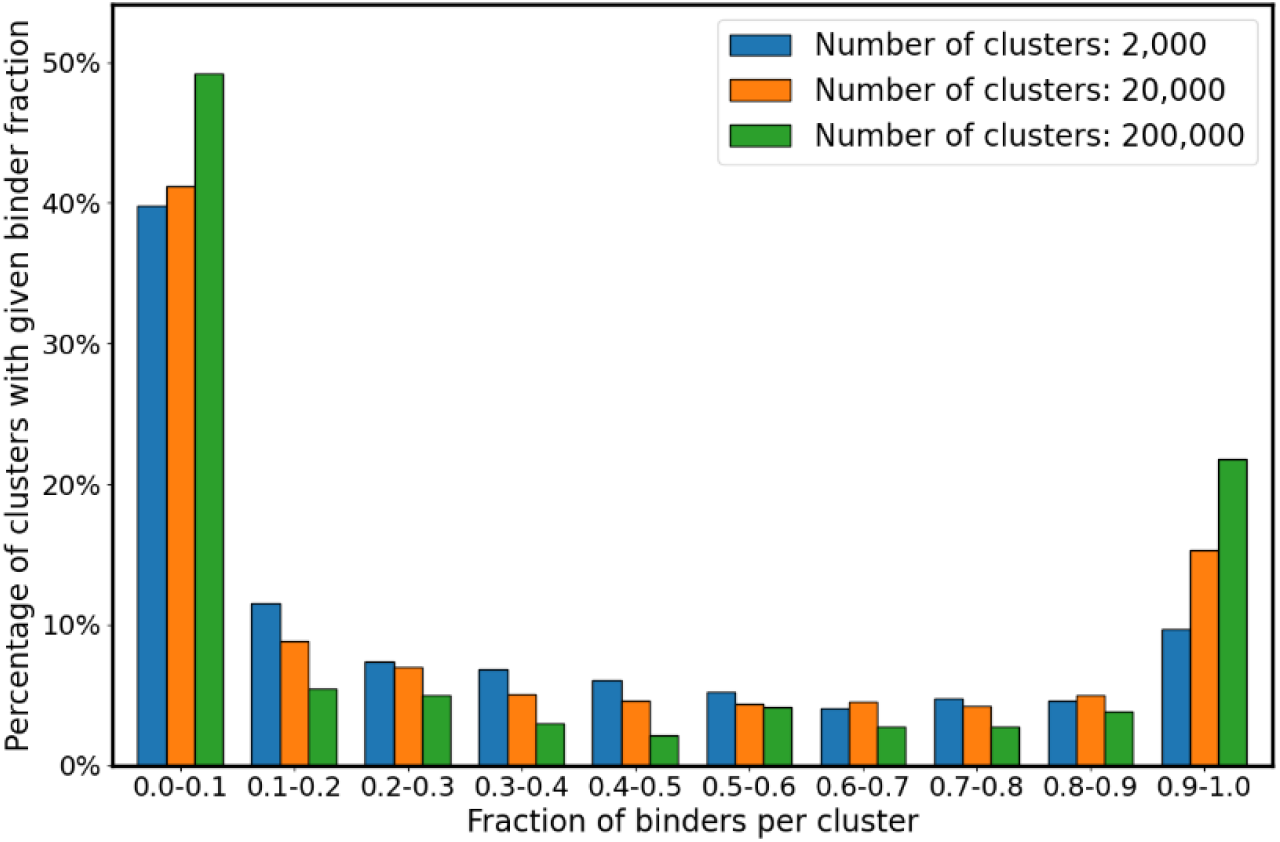
Results from k-means clustering with requested cluster sizes of 2,000, 20,000, and 200,000, illustrating predicted survivin binding fraction of peptides in the individual clusters.

Increasing the number of clusters from 2,000 to ≈200,000 provides finer resolution of potential binding behaviors; however, the proportional distribution of clusters across binder fraction ranges (e.g., 0.0–0.1, 0.4–0.5, or 0.9–1.0) remains similar. This stability suggests that increasing granularity merely divides the proteome into smaller subsets without affecting the coarse topology of binding tendencies. At lower cluster numbers, the boundary between survivin-binding and non-binding domains spans relatively more clusters. As the total number of clusters increases, these boundaries become more sharply defined, concentrating within smaller fraction of the total clusters. At the extreme limit, where each cluster contains only segments with a unique composition, the survivin-binding fraction is either 0 or 1, effectively eliminating any intermediate boundary regions.

Outside of these extremes, if the predicted composition space and its supporting protein segments were more complex, or if peptides were randomly assigned to clusters, the distinct peaks at 0.0–0.1 and 0.9–1.0 would disappear, and binder fractions would be more evenly distributed, resulting in a Gaussian histogram with a mean reflecting the overall ratio of survivin-binding to non-binding protein segments. In contrast, clustering results in two peaks, reflecting biologically relevant patterns, such as clusters with predominantly non-binding peptides or those enriched in survivin binders.

### Ontological mapping of a cluster containing solely protein segments predicted to be preferred by survivin

Next, we asked what the proteins within each cluster have in common and the list of proteins that contain compositionally similar segments was subjected to enrichment analysis by the ToppGene service ^24^ with primary focus on functional gene ontology and interactions. Gene Ontology organizes annotation by “molecular function”, “biological processes” and “cellular component”, while “interaction” groups interaction targets to an interactor protein.

Figure 4 depicts a cluster with a well-defined composition that survivin is predicted to bind. Unlike grouping proteins by a high overall fraction of survivin-binding segments (high R_bind_)^10^, it is sufficient for a protein to contain a single short segment that meets the survivin-binding criterion to be included in the enrichment query. As Figure 4a shows, the terms “chromosome centromeric region,” “mitotic cell cycle process,” “chromatin binding,” and “chromatin remodeling” are all processes in which survivin is known to be involved. What makes this cluster distinctive is its higher-than-average content of carboxyl groups (Figure 4b). Somewhat counterintuitively, the γ- and δ-carbon atoms are reduced, despite both aspartate and glutamate containing a Cγ atom. This suggests a depletion of longer aliphatic residues. Glycine Cα and guanidino groups (present in arginine residues) occur at levels typical of the proteome; however, when paired with an enriched group such as the carboxyl group, even a typical level indicates a favored presence. Composition is also strongly characterized by the groups that are completely absent such as indole (present in tryptophan residues), imidazole (present in histidine residues) and other groups.

**Figure 4.**
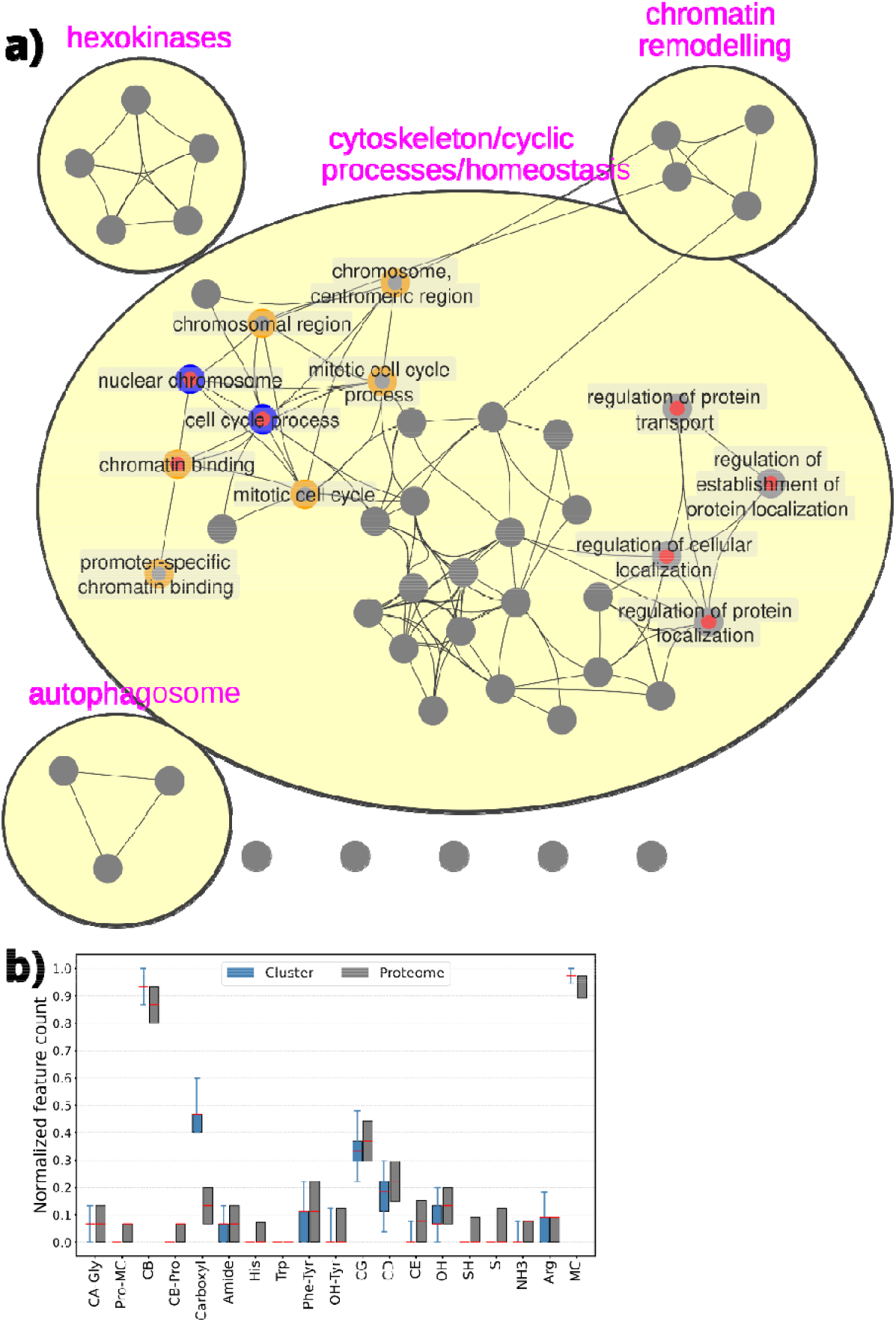
a) Functional enrichment of a cluster containing protein segments predicted to bind survivin. This cluster comprises 209 segments from 199 different proteins, but not from survivin itself. Red, orange, and blue indicate CLOCK, EZH2, and TIMELESS, respectively, in the intersection with the term. The p-value cutoff for the enrichment analyses was set at 0.05, and edges were drawn only for node intersections with a Dice similarity greater than 0.3. b) Boxplot of the normalized composition of protein segments within the cluster (blue) and within the human proteome (gray). The median is indicated by a red mark; whiskers represent the minimum and maximum of the cluster. The minimum and maximum values for proteome segments are 0 and 1, respectively.

This cluster also contains a segment of the Enhancer Of Zeste 2 Polycomb Repressive Complex 2 Subunit (EZH2, an epigenetic regulator), together with the gene products of CLOCK and TIMELESS involved in the regulation of circadian rhythms. We have shown that EZH2 as part of the Polycomb Repressive Complex 2 (PRC2) has tight functional and physical connection in the cell ^21^. CLOCK and TIMELESS are functionally coupled components of the eukaryotic circadian clock, yet they descend from entirely different protein families and share no detectable homology. Their co-occurrence in the cluster might therefore suggest functional co-evolution of protein-segment composition, possibly shared among proteins controlling functions such as the circadian clock. Survivin and the circadian clock were shown to co-regulate the chronotoxic response to cyclin-dependent kinase (CDK) inhibitors in colon epithelial cells ^25^. Survivin (BIRC5) expression follows circadian oscillation (peaking at zeitgeber time 3, ZT3) and siRNA-mediated *BIRC5* silencing selectively heightened seliciclib toxicity ^25^. It is notable that circadian rhythm disruption in hepatocellular carcinoma results in elevated survivin level worsening prognoses ^26^. Coevolution of protein-segment composition is less improbable than sequence coevolution, because only the amino-acid type of mutations matters; its exact position is of secondary importance. Conversely, a protein segment composition is sufficiently robust that single point mutations are unlikely to radically alter its functional meaning. Figures 2 and 4 show that broad compositional variation still yields similar survivin binding, a primitive protein function.

### Ontological mapping of compositional signatures related to the 106-120 region of survivin (BIRC5_106-120_)

This line of thought — namely, the possibility of identifying co-evolved proteins with similar compositions and functions — enables us to investigate which proteins resemble survivin in composition and to propose an additional hypothesis: that distinct cluster compositions may signal underlying principles of biological organization. As discussed earlier, it is well known that short signal peptides direct proteins to different cellular compartments and that these segments are typically compositionally biased ^27-29^. Here, however, we propose that the signals at the scale of 2,000-200,000 (the number of generated clusters) is relevant to biological organization.

The hypothesis is first illustrated by the interaction networks defined by the proteins that contain a 15mer segment resembling 106-120 segment of survivin. Figure 5a presents the enrichment analysis of GO terms for proteins within one of the 20,000 clusters containing BIRC5_106-120_. Proteins with segments similar to BIRC5_106-120_ are more likely to be involved in microtubule binding, spindle assembly, chromosome condensation and segregation, cell division, and methylated histone binding—all known functional domains of survivin. Some associated terms directly include survivin, while others, even in its absence, are linked to its functions. For example, the term “chromatin” does not explicitly include survivin, despite growing evidence of its chromatin-associated roles. Other related proteins do not co-occur across all terms. For example, Shugoshin-2 (SGO2) and survivin exhibit substantial overlap, as SGO2 is known to be involved in sister chromatid cohesion, and survivin interacts with shugoshins ^30-32^. In contrast, Rho-associated protein kinase 1 (ROCK1) occupies a distinct domain within this functional network, with less overlap, particularly in centriole- and microtubule-related functions. ROCK1 and survivin are not known to bind to each other directly. Despite their functional associations, survivin, SGO2, ROCK1, and other proteins in the composition-based cluster do not share sequence homology. We associate them solely based on a shared small segment with similar functional group composition, defined by 19 parameters of C-Pos encoding. Despite the simplicity of the model, the accuracy and precision of functional assignment are far from vague (e.g., generic terms like “cytoplasm” or “protein binding”). Instead, the assignments are highly specific and relevant, targeting distinct cellular structures (condensed chromosome centromeric region due NCAPD3, ATRX, CENPF, BIRC5, GPATCH11, SGO2) and precise functions (sister chromatid segregation due to NCAPD3, GOLGA8J, CENPF, GOLGA8K, BIRC5, AKAP8L, GOLGA8M and GOLGA8H).

**Figure 5.**
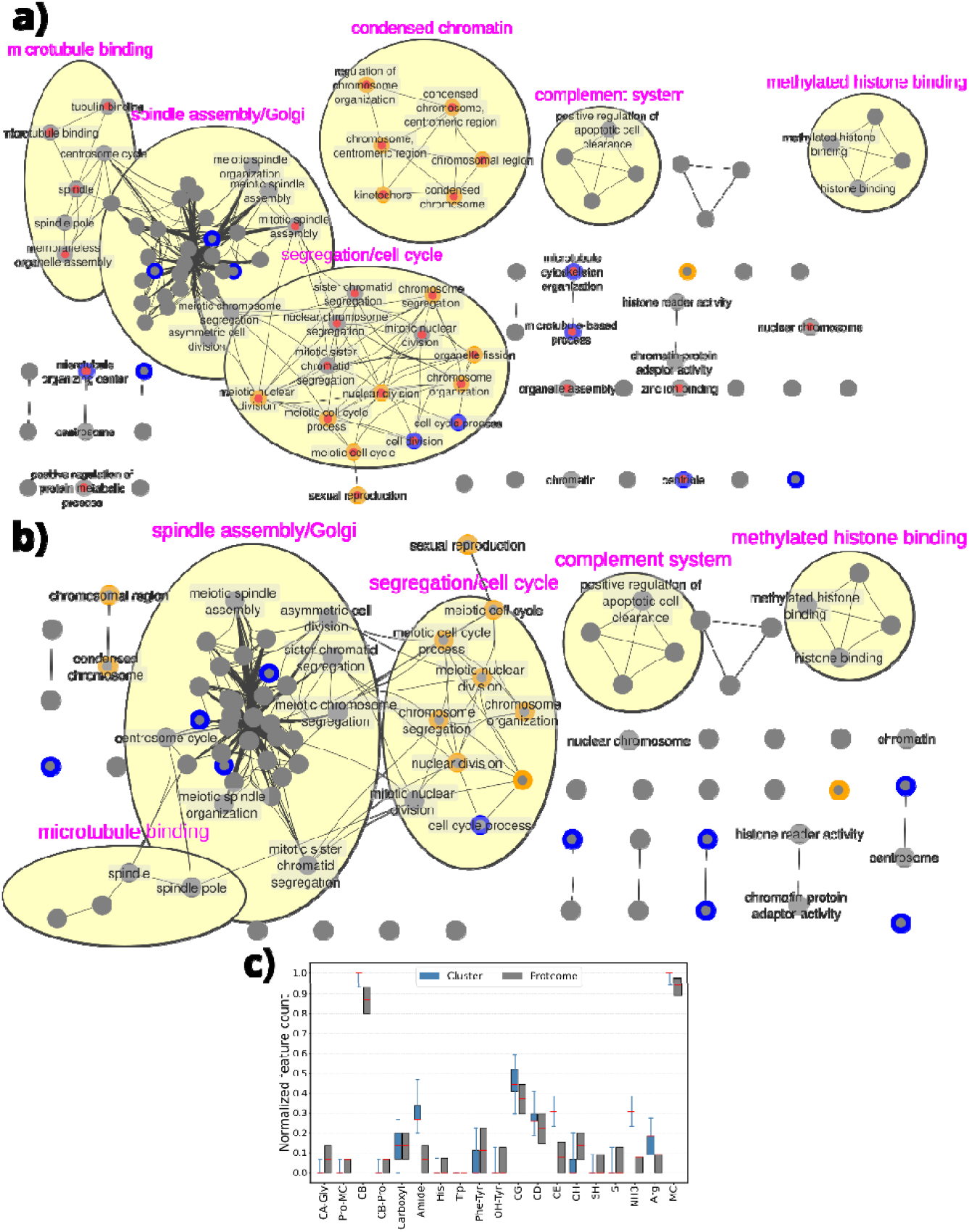
a) A densely connected functional enrichment of proteins in the cluster that includes BIRC5_106-120_. This includes 133 protein segments that belong to 124 proteins. 94% of these protein segments are predicted to bind survivin. Red, orange and blue indicate survivin, SGO2 and ROCK1 in the intersection with the term, respectively. b) The same cluster without including survivin in the enrichment analysis. The p-value cutoff was set at 0.05 for both enrichment analyses, and edges were drawn only for node intersections with a Dice similarity greater than 0.5. c) Boxplot of the normalized composition of protein segments within the cluster (blue) and within the human proteome (gray). The median is indicated by a red mark; whiskers represent the minimum and maximum of the cluster. The minimum and maximum values for proteome segments are 0 and 1, respectively.

The presence of survivin was a factor in selecting this cluster, so a bias toward survivin-related functions is expected. However, when survivin was excluded from the list of proteins subjected to enrichment analysis, the overall topology of enriched functions remained largely unchanged (Figure 5b). This suggests that the bias introduced by survivin is minimal and that the identified functions are robustly described by the collection of proteins within the composition-based cluster. The nature of the compositional signal is illustrated in Figure 5c. The most notable feature of this cluster is the increased amide, amino and Cγ content in the protein segments. Guanidino groups are also somewhat elevated. Glycine, proline residues are almost completely absent.

When using ≈200,000 (Figure S6) or 2,000 (Figure S7) total clusters, the enrichment analysis has yielded additional insights. One might expect that low granularity in clustering—grouping an excessive number of proteins into broad clusters—reduces specificity, while overly fine-grained clustering may result in sparse protein representation, introducing artificial divisions even among highly similar peptides and thereby complicating enrichment detection. In Figure S6a, a prominent survivin relevant term is “cytokinesis” which it performs together with ROCK1. An example of a fine-grained insight is that both survivin and ROCK1 are targeted by apoptotic proteases (caspases) ^33,34^ and they get completed by Serine Protease Inhibitor Kazal-type 5 protein (SPINK5) which is an anti-inflammatory protein that inhibits KLK5, KLK7, KLK14, CASP14 ^35^, and trypsin ^36^. In Figure S7b, the elevated levels of amide, amino and Cε groups is apparent. Functional networks associated with proteins that share segments with this composition include highly interconnected and broad cellular functions. Survivin and SGO2 are recognized mostly by their association with mitotic chromosome segregation and associated functions, but survivin is also recognized with its association with cytoskeleton and centriole together with ROCK1. As an example “microtubule organizing center” association is due to MCM3, JADE1, SGO1, CCDC112, MTUS1, STARD9, CCDC15, NLRC5, CAMSAP1, SLAIN2, FLII, ATF5, CEP162, CEP290, MYO18A, BIRC5, CCDC77, CCDC88A, NINL, TNKS, ATF4, RAB11FIP4, INTU, SPAST, TTLL9, NUBP1, CFAP100, ABRAXAS2, CCDC66, ARHGEF7, NFE2L2, KIF2C, TCEA2, TBC1D30, CAPG, GSK3B, KIF3A, ORC2, CEP89, ERCC6L2, PCM1, PDE4D, CHD3, CHD4, RABGAP1, KIF3B, RRAGD, ASPM, EZR, RNF19A, KIF20B, TUBGCP6, PPP2R5A, C2CD3, ITSN2, PRKAR2A, PRKAR2B, PRKCB, PKN2, MAP2K1, CCDC65, CCP110, AHI1, KIAA0586, HAUS3, PTK2, CEP350, EVI5, ALMS1, KIF2A, DYNC1H1, LCA5, CCSAP, C10orf90, TTLL7, AKAP9, CEP83, LRP1, CFAP58, ROCK1, TXNDC9 and TOPORS contained in this cluster. Many other familiar gene products appear which contribute to survivin related functions in different contexts such as Inner Centromere Protein (INCENP) ^37^, Shugoshin-1/2 (SGO1/SGO2) ^31^ and SUZ12 ^21^.

### Protein association networks associated with composition of BIRC5_106-120_

Next, we examined the experimentally verified interactions reported in the STRING database between the proteins that belong to this cluster. In Figure 6a, the k=20,000 cluster includes a tightly interwoven network of interacting proteins, as indicated by the STRING database, suggesting that similar signaling fragments, based on composition, exchange information through binding events. We note that BIRC5 is linked to SGO2 through Centromere protein F (CENPF). In the likely incomplete physical subnet BIRC5 is only associated directly with CENPF (Figure 6b). In the STRING network of fine-grained k=200,000 cluster (Figure 6c) BIRC5 is more closely associated with ROCK1 with Heat Shock Protein 90 Alpha Family Class A Member 1 (HSP90AA1) taking a central place. We will see that Heat Shock Protein 90 Alpha Family Class B Member 1 (HSP90AB1) also displays compositional similarity to BIRC5. This is notable given their distinct primary functions—HSP90 as a molecular chaperone and BIRC5 survivin as an apoptosis inhibitor and mitotic regulator—yet their local segments suggest convergent composition. Importantly, the functional interaction between HSP90 and survivin has been extensively studied over the past two decades. Survivin is stabilized by forming a complex with HSP90, protecting it from proteasomal degradation and enabling its role in mitosis and cytoprotection. This interaction has been therapeutically targeted by shepherdin, a cell-permeable peptidomimetic derived from the survivin–HSP90 interface, which disrupts their complex, leading to tumor-selective apoptosis and growth suppression ^38,39^.

**Figure 6.**
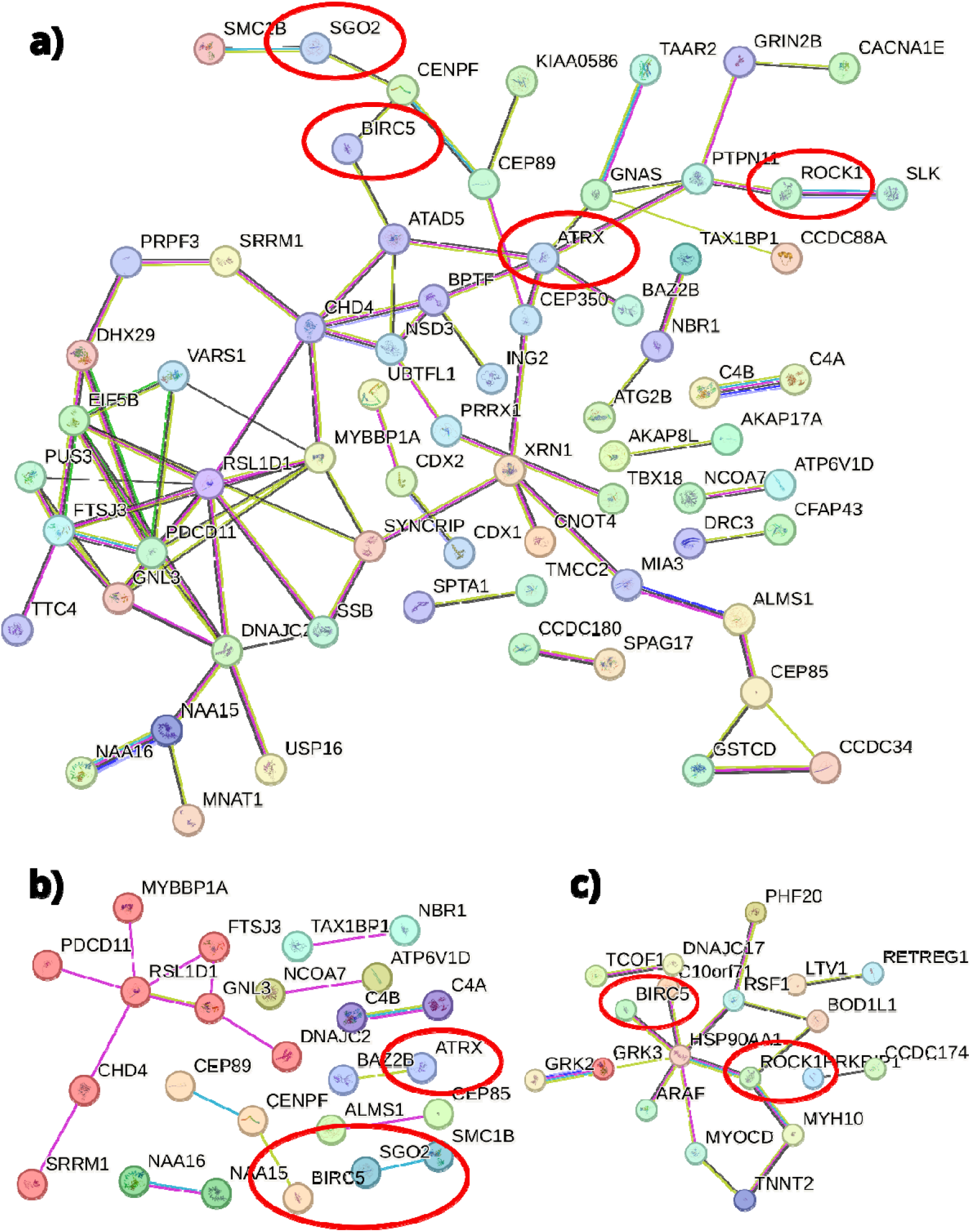
STRING network analysis. a) Full STRING network and b) physical subnetwork for a k=20,000 cluster that includes BIRC5_106-120_. c) Full STRING network for a k=200,000 cluster that includes BIRC5_106-120_.

Transcriptional regulator ATRX (when present in Figure 6a) is also one step away from BIRC5 and it is associated with the centromeric region. Interestingly, ATRX, a helicase belonging to the Switch/Sucrose Non-Fermentable (SWI/SNF) family ^40^, also binds to EZH2 ^41^ of the PRC2 complex. Both the SWI/SNF and PRC2 complexes have been shown to associate with survivin as well ^21,42^. The association between these proteins is likely attributable to their shared signaling composition within the fragment, suggesting that compositional similarity is a property that facilitates their interaction.

### Compositional signatures related to the upstream 101-115 region of survivin (BIRC5_101-115_)

This composition is characterized by a relatively high amino group content and increased Cδ and Cε content, primarily due to lysine rather than arginine. Indole and thioether (present in methionine residues) groups are entirely absent, and hydroxyl groups are reduced in presence. In contrast, amide and carboxyl groups are moderately elevated.

The most prominent functional association is observed in Figure S8, in a tightly interconnected subgroup of proteins involved in bypassing the spindle assembly checkpoint and facilitating the metaphase-to-anaphase transition during cell division. These functionally related proteins form a nearly fully connected network, despite the stringent intersection similarity requirements used in the analysis. Survivin plays a central role in this group, particularly in the highly specific negative regulation of spindle checkpoint activity. The full list of these functionally associated proteins includes: KNTC1, DLGAP5, NUP62, TRIP13, Aurora kinase A (AURKA), MKI67, ZWILCH, BIRC5, CDK5RAP2, GEN1, PBRM1, and SMARCC1. Still connected to these functional network survivin dominated cluster with examples of “mitotic cell cycle”, “spindle assembly” and “chromosome segregation”. Here ATRX has an increasing importance in addition to AURKA and BIRC5. Chromosome segregation is facilitated by KNTC1, USP9X, SMC1A, DLGAP5, NUP62, TRIP13, GOLGA8N, AURKA, SYCP2, TUBGCP3, MKI67, KIF14, ZWILCH, KAT5, KNSTRN, BIRC5, HAUS2, PPP1R7, GOLGA8M, PBRM1, SMARCC1, SASS6, CDK5RAP2, GOLGA8O, GEN1 and RNF212B within this cluster.

This region is part of the dimerization interface of survivin as F101 and L102 forms contact between the subunits. Aside from survivin, several other homodimerizing proteins feature in this cluster, including LRRK2, RNF40, BLM, DST, PIP4K2A, TRIM8, PIP4K2B, TAP1, HSP90AB1, RAG1, KNSTRN, SYNE1, SMCHD1, CGAS, WRN, MSH3, CRYL1, TRNT1, RABEP1, PYCARD, APPL1, GEN1, and TP53BP2. It raises the possibility that this compositional signature promotes homodimerization.

The STRING network in Figure S8b is extensive, but BIRC5 is only connected to AURKA. ATRX is part of a network in which HSP90AB1 and Epidermal Growth Factor Receptor (EGFR) occupy central positions. SMARCC1, a component of the SWI/SNF chromatin remodeling complex, is linked to this network through histone acetyltransferase KAT2A, highlighting an extended axis of chromatin regulation and signal transduction.

The segments 101–115 and 106–120 overlap by 10 amino acids and are therefore not independent; however, subtle compositional differences place them in distinct clusters. If other proteins repeatedly co-occur with both segments, it likely reflects distinct regions within those proteins, as a given segment can only belong to a single cluster. To describe such co-occurrence of segments across two or more proteins, we use the term **“compositional segment intersection**,**” (CSI)** referring to either point-like or continuous similarities in segment composition.

### Survivin has a well populated CSI with homogenous composition (BIRC5_111-125,_ BIRC5_116-130_)

The functional enrichment of this region describes many known survivin functions such as cytokinesis which includes microtubule binding, spindle and chromosome organization and the act of cytokinesis itself. AURKA is part of this cluster as well and we selected Structural Maintenance of Chromosomes 4 (SMC4) to illustrate a partially overlapping set of functions that BIRC5, AURKA and the condensin ^43^ complexes have in Figure S9. In the STRING network, these three proteins are centrally located, recognizing that BIRC5 and AURKA is part of the Chromosome Passenger Complex and Aurora kinases are known to change the composition of condensin by phosphorylation^44,45^.

BIRC6 (BRUCE/APOLLON) and BIRC5 both contain a BIR domain, so their compositional similarity might first be ascribed to this shared motif. In fact, the segments under comparison do not arise from the BIR domain: in BIRC5 the segment lies in its unique C-terminal helix, whereas in BIRC6 it originates from a ubiquitin-like domain. This rules out straightforward homology and instead points to compositional co-evolution due to shared (anti-apoptotic) function as the more plausible explanation. HSP90AA1 reappears as a centrally positioned hub in the STRING network directly linked to BIRC5 (Figure S9b).

We note the presence of clathrin related functions that were previously proposed based on clathrin components with high R_bind_ values ^10^ with gene products SMAP1, COPB1, MYCBPAP, AP1B1 and AP2B1. The functional network includes “chromatin remodeling” due to TTF2, SETD1A, ARID4A, KDM5A, BAZ2B, SMCHD1, CHD2, BRD3, USP16, AURKA, BMI1, SETD2, UBN1, BAZ2A, CTR9 and ATAD2B.

When examining the nature of the compositional signal, amino and Cε groups stand out, indicating an elevated presence of lysines. Amide and carboxyl groups are also enriched. Using the examples BIRC5_116-130_ (NKIAKETNNKKKEFE), AP1AR_81-95_ (EKQKDLDKKIQKEL) and BIRC6_3881-3895_ (AKLISEQKDDKEKKN), we illustrate that permutations of either identical or different residues can yield similar overall chemical group composition. A clear example is the N–E and D–Q residue pairs, which are equivalent in C-Pos and many other compositional encodings. In BIRC5_116-130_, only N and E are present, whereas AP1AR_81-95_ is rich in D and Q. However, these letter differences are superficial and inconsequential in terms of chemical group composition.

### Preferred and disregarded compositional signatures in the human proteome

Beyond this selected example, we also investigated how many clusters could be associated with enriched *interaction targets*. To address this, we performed a ToppGene enrichment analysis of “Interactions,” which links an *interactor* to multiple *targets*. An *interactor* may frequently serve as its own *target* if it is known to self-associate into higher-order assemblies, though this is not always the case. Figure 7a shows that approximately 8,000 of the 20,000 clusters do not have any enriched interaction targets to any of the known interactors. But, more than half of the assessed clusters have at least one associated *interactor*, and 2,000 clusters contain 100 or more associated *interactors*. This suggests that thousands of compositional signals are present in the proteome, with a substantial fraction of these robust signals being recognized or targeted by more than 100 interactors.

**Figure 7.**
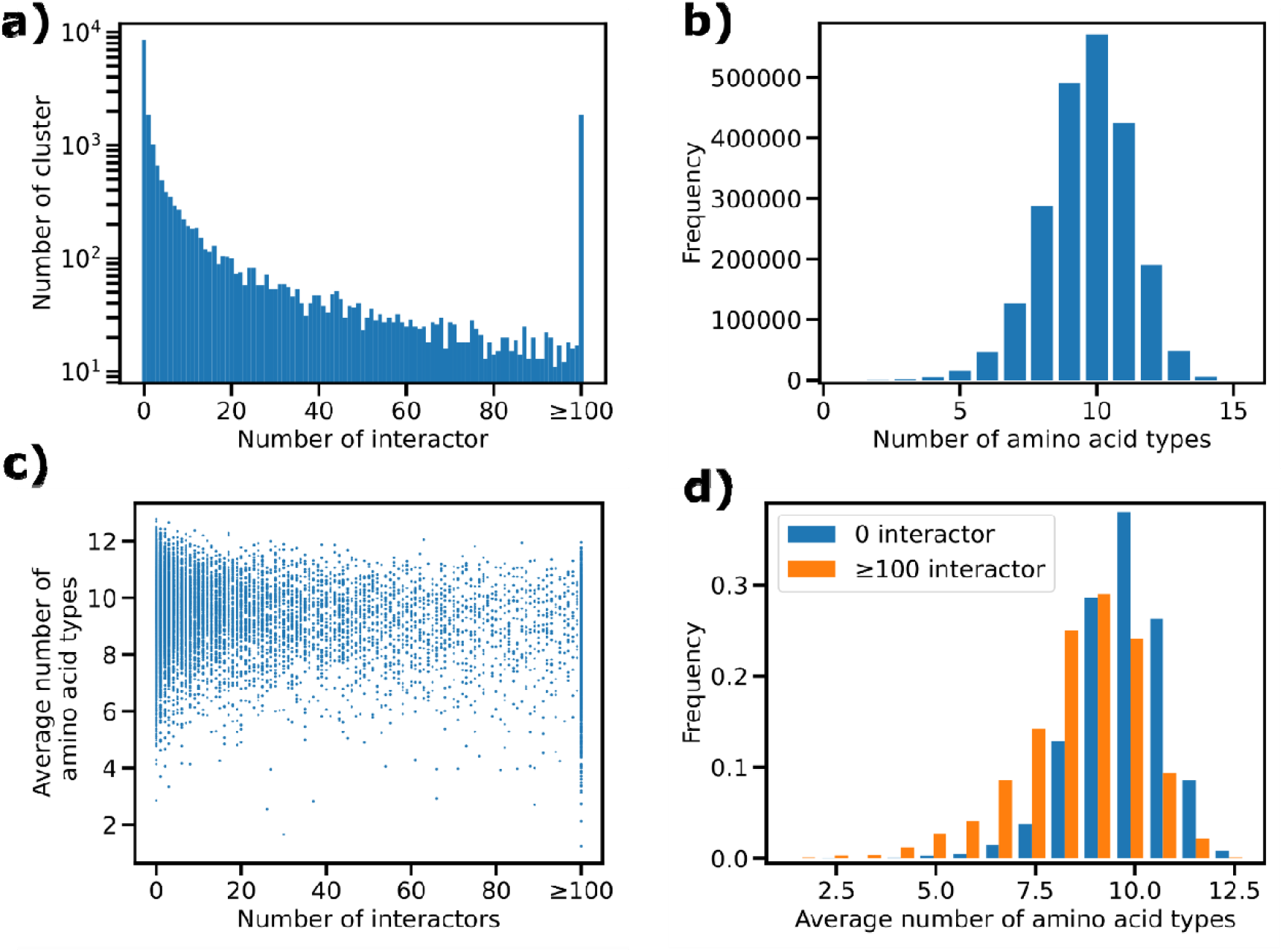
a) Number of interactor proteins associated with the peptide clusters, as determined by ToppGene enrichment. The maximum number of interactors was limited to 100. b) Histogram depicting the distribution of the number of amino acid types in 15-mer protein segments from the human proteome. c) Scatterplot showing the number of interactors versus the average number of amino acid types in cluster peptides. Each dot represents a cluster. d) Average number of amino acid types in clusters with either 0 or more than 100 associated interactors with frequencies normalized to represent probability densities.

Next, we investigated how protein segment complexity/purity influences their ability to serve as strong signals, i.e., having many interaction partners. We analyzed the number of different amino acid types in 15-mer peptides derived from the human proteome (Figure 7b). The most common 15-mer segment contains 10 different amino acid types. Segments composed of a single amino acid type are rare but occur at orders of magnitude more frequently than expected from random sequences. Similarly, segments with more than 10 different amino acid types become progressively less common, aligning with probabilistic expectations.

The average number of distinct amino acid types in peptides across the 20,000 clusters shows a weak correlation (Pearson CC −0.28) with the number of associated interactors (Figure 7c), suggesting that the two variables are not fully independent. Figure 7d compares the complexity (average number of different amino acid types) of clusters with 0 enriched interactors to those with more than 100 interactors. The distribution for clusters with over 100 interactors is shifted toward fewer amino acid types, but compositions with typical complexity/purity can also identify proteins with many common interaction partners. It is important to highlight the relationship between a small segment of a protein and the number of binding partners of the protein. If a small segment were not influential in determining the binding partners of a protein, the histograms in Figure 7d would appear more similar, and other properties of the whole protein, such as its overall structure, shape and surface, would only be required to determine the number of interactors.

### Limitation of this study

It is important to mention the limitations of our clustering and enrichment analysis. First, although we have explored a few encodings other than C-Pos previously ^10^, there are still other alternatives that might align better with the physical processes underpinning protein-protein interactions. Second, the exact choices of 2,000, 20,000 and 200,000 clusters remain somewhat arbitrary, and they may not precisely capture qualitative compositional signals. It is important to note that K-means clustering is non-hierarchical; therefore, fine-grained clusters do not necessarily align with their coarser counterparts. The degree to which the survivin segment features deviate from the cluster centroid (i.e., how representative the cluster composition is for the survivin segment) also varies between clustering attempts. At the lowest granularity, the variation in compositions is understandably greater, yet remains specific enough when compared to the 0–1 scale of the normalized proteome distribution. Third, the segment window of 15 amino acids is also an arbitrary choice, aligned with the peptide microarray design. However, the optimal segment size requirement for a compositional signal remains unclear, and it is uncertain whether such an optimum even exists. Combinatorial arguments suggest that a minimum length is necessary to represent enough signals. Fourth, enrichment analysis is sensitive to the typical sizes of the query and terms, with certain combinations inherently unsuitable for statistical enrichment a priori. Fifth, the clustering analysis unintentionally enriches for proteins with longer sequences, complicating accurate modeling of the resulting biases. These proteins tend to accumulate because they are represented by a greater number of segments, and the clusters become crowded with segments derived from longer proteins. While longer proteins may indeed engage in more interactions and functions if they possess sufficiently diverse properties, assuming that these properties are solely compositional may be misleading. Consequently, GO terms associated with long proteins are particularly susceptible to enrichment due to length bias.

The same arguments apply to grouping proteins by other extensive properties (for example, surface diversity in terms of distinguishable hydrophobicity, charge, or hydrogen-bonding patterns, as well as domain or sequence motif diversity). This underscores the need for caution when interpreting hypotheses derived from enrichment analyses. The only reason our features are more compelling than others is that they can be effectively interpreted using extremely simple models, which are validated by the microarray experiment described here.

### Conclusions

Our findings suggest a robust and parsimonious framework for decoding protein interactions based solely on compositional features. In Figure 8, a protein is represented as a continuous curve in a 19-dimensional compositional space, generated by sliding a segment window from the N-to the C-terminus. Self-similarity and chain-like continuity in this space are maintained by the overlap between adjacent windows—a property that is especially pronounced when the window advances by a single amino acid, rather than by the five-residue step used in our study.

**Figure 8.**
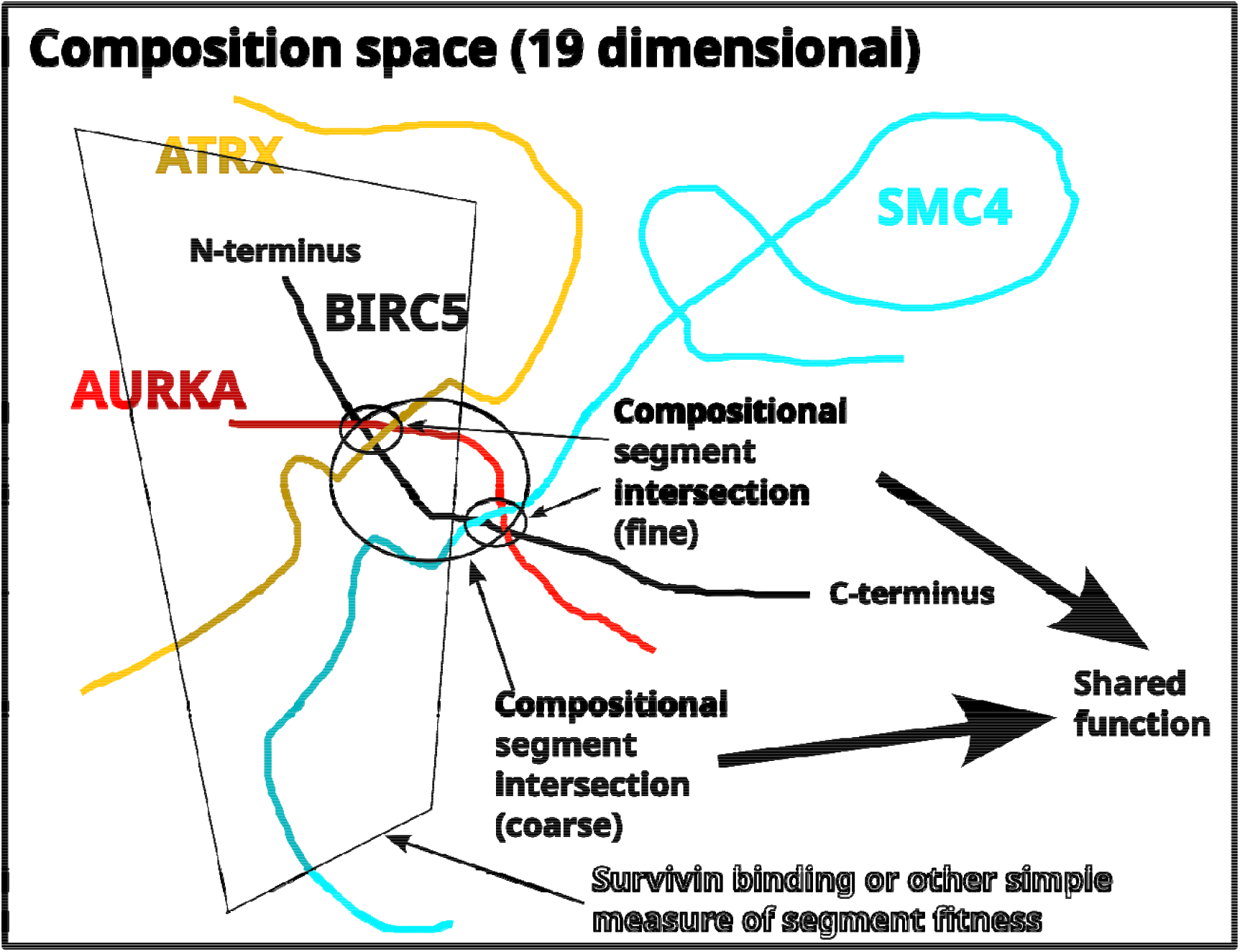
Conceptual visualization of polypeptide chains in a composition space. Using different similarity criteria, both fine- and coarse-grained compositional intersections are defined. The uncolored regions represent compositional space not occupied by the human proteome. The shaded gray area denotes a continuous region of compositional space that meets a fitness criterion, such as the potential to bind survivin.

Compositional space is not astronomically large; consequently, segments from different proteins may occur in close proximity to one another—whether by chance, sequence duplication, or compositional co-evolution. Depending on the level of similarity accepted, one can define finer or coarser **compositional segment intersections**. We hypothesize that these intersections represent signals of a coherent set of functions shared by the participating proteins. From a physico-chemical perspective, such segments may often correspond to imperfect mixtures with an unfavorable free energy of mixing, as described by a variant of the Edmond–Ogston model. Since chemical groups of the side chains cannot migrate along the polypeptide chain to approach an energy minimum, the only route toward a more ideal mixture is through association with other proteins or compatible compounds. If a protein segment is relatively close to an ideal mixture, it may associate with another segment that is similar yet complementary. For illustration, consider the hypothetical case in which a 1:1 mixture of **S** and **Y** represents an ideal mixture. Associations between segments such as **SYSYYYSYSY** and **SYSYSSSYSY** approach ideality and can still be detected within the same cluster due to inherent similarity. Such segments and their complements can be identified in a CSI, with detection becoming more likely as the clustering is made more coarse-grained. Favorable association may also occur between highly dissimilar low-complexity regions, in which case an accurate description of the virial terms becomes essential and the mechanisms underlying their values must be understood. An example is given by the pure segments **YYYYYYYYYY** and **SSSSSSSSSS**, which, although yielding the same net composition, cannot have their complementarity detected on the basis of similarity.

Proteins in composition space may intersect multiple times, or align in parallel or antiparallel— either with one another or with themselves—emphasizing certain functional features. We have also observed that survivin exhibits considerable tolerance to compositional variation when recognizing binding peptides. This functional plasticity must be taken into account when defining the scope and thresholds of relevant intersections. Importantly, this representation does not require continuity of the polypeptide backbone across the entire protein. Dividing a protein into two or more larger fragments does not alter the compositional space these segments occupy— apart from the introduction of new termini. This suggests functional independence of distinct protein regions and supports the notion of transferable biological properties in chimeric constructs or fragment cocktails, thereby de-emphasizing protein chains as functional units in biology.

By reducing the complexity of protein-protein interaction prediction to a 19-dimensional compositional space, we demonstrate that functional group composition captures biologically meaningful signal for survivin binding, independent of sequence or structural homology. The predictive accuracy of this model, validated across diverse experimental datasets, reveals that protein interactions can be governed by simple compositional logic. Clustering across the human proteome hints towards a latent compositional architecture aligned with functional networks, suggesting that biological specificity and higher-order organization may emerge from a limited repertoire of recurring compositional motifs. This new hypothesis—comprising three layers: protein chains, clusters of compositionally similar protein segments, and the functional enrichment of proteins within these clusters—is unified by the composition of short segments.

Viewed in this broader context, the framework offers insight into the evolution of molecular complexity. By describing proteins and their interactions through compositional similarity rather than sequence ancestry, it suggests that early protein–protein associations may have emerged from simple physico-chemical compatibilities among primitive segments. This view links modern protein networks to compositional self-organization processes that likely preceded the advent of accurate sequence-based heredity, which employs the ribosome to print precisely defined mixtures of chemical groups. The evolution of interaction specificity may thus reflect a gradual refinement of ancient compositional constraints. Revisiting current models of protein and PPI evolution from this perspective highlights the broader significance of compositional logic in shaping the molecular architecture of life.

## Methods

### Peptide microarray experiments and data reduction

Protein sequences were elongated at both N- and C-termini with neutral GSGSGSG linkers to prevent truncation. These extended sequences were segmented into linear 15-mer peptides with 10-amino-acid overlaps, yielding a peptide microarray comprising 5,376 unique peptides, each printed in duplicate (10,752 spots). The array was framed with 232 HA control peptides (YPYDVPDYAG). Prior to the main assays, the peptide microarray was pre-stained for 45 minutes with secondary (Mouse anti-6x-His tag [HIS.H8], DyLight680, 0.5□µg/mL) and control (Mouse monoclonal anti-HA [12CA5], DyLight800, 0.2□µg/mL) antibodies to assess nonspecific background interactions.

A separate peptide microarray was incubated with His-tagged survivin at concentrations of 10□µg/mL, followed by staining with secondary and control antibodies and subsequent microarray read-out. The HA peptides framing the array were concurrently stained with the control antibody, serving as internal quality controls to validate assay performance and array integrity.

Rockland Blocking Buffer MB-070 was used for blocking (30□min prior to the first assay). Washing steps were carried out using PBS (pH 7.4) containing 0.005% Tween 20, with two 10-second washes after each incubation. The incubation buffer consisted of 90% washing buffer and 10% blocking buffer.

Microarrays were scanned using an Innopsys InnoScan 710-IR Microarray Scanner at 20□µm resolution. Scanning parameters were set to a gain of 50 at low laser power (680□nm, red channel) and 10 at high laser power (800□nm, green channel). Spot intensities and peptide annotations were quantified from 16-bit grayscale TIFF files. Image analysis was performed using PepSlide® Analyzer and results were compiled in Excel. The software extracted raw, foreground, and background signals for each spot, calculated averaged median foreground intensities, and assessed spot-to-spot variability between duplicates. A maximum spot-to-spot deviation of 40% was tolerated; higher deviations resulted in the corresponding intensity value being set to zero.

### Machine learning methods

To evaluate the neural network models across the full proteome, we trained our model on a dataset previously published^21,46^, which contained sequence fragments from the following proteins: Cdk1 (P06493), KAT2A/GCN5 (Q92830), SPI1/PU1 (P17947), SUZ12 (Q15022), EED (O75530), JADE3 (Q92613), DIABLO/SMAC (Q9NR28), BOREALIN (Q53HL2), INCENP (Q9NQS7), SGOL1 (Q5FBB7), SGOL2 (Q562F6), EZH2 (Q15910), JARID2 (Q92833), Histone H3 (P68431), AURORAKB (Q96GD4), JADE1 (Q6IE81), JTB (O76095), EVI5 (O60447), RAN (P62826), USP9X (Q93008), C-IAP1 (Q13490), STAT3 (P40763), BRUCE/APOLLON (Q9NR09), XPO1 (O14980), CDX2 (Q99626), Msx2 (P35548), RBM15 (Q96T37), PHF21A (Q96BD5), PHF8 (Q9UPP1), DIDO (Q9BTC0), JADE2 (Q9NQC1) and HASPIN (Q8TF76).

We used a multilayer perceptron neural network implemented in scikit-learn with default parameters, except for the model consisted of a single hidden layer with only 2 neurons. Despite its simplicity, this model was found to outperform more complex alternatives^10^ and was therefore selected for all subsequent analyses. The features were generated by the C-Pos encoding^10^ which categorize non-hydrogen atoms of each amino acids according to 19 categories: main chain atoms (**MC**), except glycine (3 non-hydrogen atoms) and proline (2 non-hydrogen atoms). Additionally, specific features included the glycine Cα atom (**CA-Gly**), recognizing its methylene group nature instead of a methanetriyl-group. Proline-specific main chain atoms (**Pro-MC**) were considered due to the cyclic nature of the proline side chain, which links the amide nitrogen to the Cα atom. The content of side chains were deconstructed to carboxyl groups of aspartate and glutamate (**carboxyl**), the amide groups of asparagine and glutamine (**amide**), the imidazole ring of histidine (**His**), the indole ring of tryptophan (**Trp**), the phenyl group of phenylalanine and tyrosine (**Phe-Tyr**), the phenolic hydroxyl group of tyrosine (**OH-Tyr**), the hydroxyl group of serine and threonine amino acids (**OH**), the sulfhydryl group or oxidized variants of cysteine (**SH**), the thioether of methionine (**S**), the amino group of lysine (**NH3**), and the guanidino group of arginine (**Arg**), the aliphatic carbon atoms were categorized by their distance from the main chain (**CB, CG, CD** and **CE**). The Cβ atoms of prolines were historically assigned a separate category (**CB-Pro**); however, as these correlate perfectly with the **Pro-MC** category, maintaining a distinct category is not justified in this implementation. The standard scaler was fitted on the entire previous peptide microarray data prior training ^21,46^ and the entire data set was used for training. The same scaling was applied to the new peptide microarray data prior to prediction (i.e., the mean and variance of the training data were used as references), ensuring the logical expectation that identical compositions are represented by the same standardized values, regardless of whether they belong to the training or prediction set.

To evaluate the performance of the model, we compared predicted and experimentally observed binding fractions, both defined as the proportion of binding peptides relative to the total number of peptides within each protein. The predicted binding fraction was recalculated as:

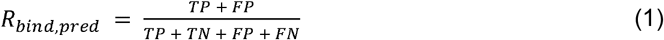

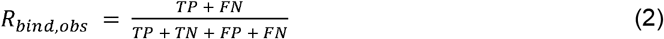

This reflects the proportion of peptides classified by the model as binders. In this context, true positives (TP) are peptides correctly predicted as binders, false positives (FP) are non-binding peptides incorrectly predicted as binders, true negatives (TN) are peptides correctly predicted as non-binders, and false negatives (FN) are binding peptides incorrectly predicted as non-binders. A *priori* the separation between the true and false fractions of categories was not known.

We further evaluated the model by calculating the accuracy for each protein, defined as the proportion of correctly classified peptides. The accuracy was calculated as:

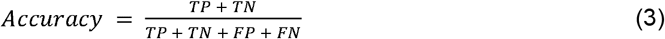

To avoid potential biases introduced by a specific training set and the model trained upon this data, we also evaluated the features directly by discarding the trained model and analyzing the feature space for clustering patterns. Here, we standardized the features and applied mini-batch K-means clustering (using default parameters from scikit-learn, version 1.6.1). With parameter k=2,000, 2,000 clusters were generated; however, k=20,000 resulted in 19,273 clusters due to mergers, and k=200,000 yielded 198,024 clusters.

The biological relevance of the resulting clusters was evaluated by identifying known interacting proteins within each cluster and conducting enrichment analysis via an in-house Python script querying ToppGene. Interaction networks based on the identified clusters were visualized using Cytoscape (version 3.10.1).

### Thermodynamic model, Bayesian inference and prediction

As an attempt to quantify the survivin binding trend, we have modeled the binding free energy of survivin to peptides based on their amino acid composition. For this purpose, a modified version of the Edmond–Ogston model (EOM) ^22^ is adopted.

The EOM is a mean-field approach used to describe liquid–liquid phase separation in mixtures of macromolecules. It relies on a truncated virial expansion of the Helmholtz free energy, retaining terms up to second order in macromolecular concentrations. Under aqueous microarray conditions (T ≈ 295 K = const, ambient pressure, nearly incompressible solutions), changes in molar volume are small (typically on the order of tens of ml mol^-1^), corresponding to a work variation of only a few J/mol. Thus, a Helmholtz free energy contribution for survivin– peptide bonds can, to a good approximation, be treated as the corresponding Gibbs free energy (*G*) change.

For a ternary mixture in a common solvent, Δ*G* can be expressed similarly to how we model Δ*G*_*bind*_ for the peptide–survivin association reaction:

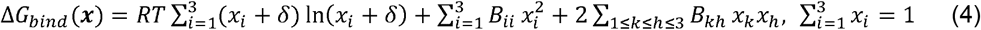

where vector *x = (x*_*1*_, *x*_*2*_, *x*_*3*_) represents the relative mole fractions of amino acids in the peptides, *B*_*ii*_ [kJ mol^-1^] are the self⍰virial coefficients, *B*_*kh*_ [kJ mol^-1^] are cross-virial coefficients and *R* is the universal gas constant in units of kJ mol^-1^ K^-1^. The formal definition of these coefficients is well-known ^22^, and entails averaging the field forces among the involved components (*ii, kh*). A small constant (δ = 10−□) was added to the amino acid content in the entropic term for computational convenience, ensuring that the absence of specific amino acids in the relevant compositions could be handled while yielding only a negligible contribution to Δ*G*_*bind*_(*x*).

When Eq. (4) is regarded at equilibrium, the van’t Hoff equation determines an effective dissociation constant for the following chemical thermodynamics scheme, involving the peptide and survivin:

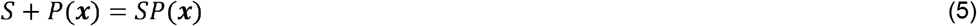

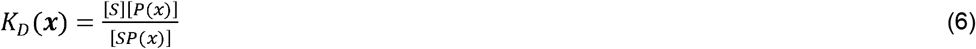

where [*S*] is the unbound survivin concentration (total c_surv_ = [*S*] *+* [*SP*(*x*)] = 6.1 × 10^-7^ M) and [*P*(*x*)] is the concentration of the unbound peptide (total concentration c_pept_ = [*P*(*x*)] *+* [*SP*(*x*)] ≡1 mM), with composition ***x***:

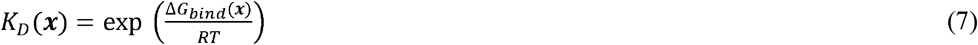

We considered that the fraction of survivin bound is governed by the law of mass action:

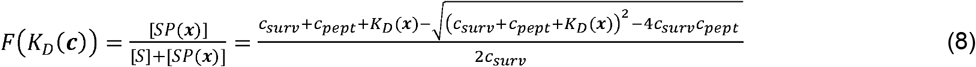

Concerning the fluorescence data (*I(****x***)), we assumed the fluorescence value is linearly proportional to the concentration of the bound survivin fraction, the maximum instrument readout (*I*_*max*_) representing the complete binding state:

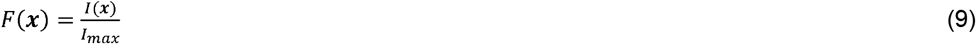

The Bayesian model incorporated the following uniform prior expectations on every virial coefficients:

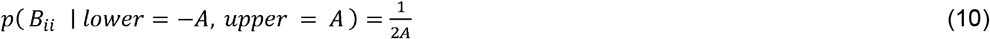

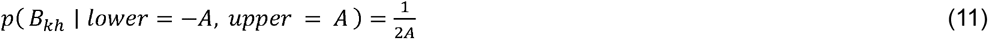

where A = 60 and A = 20 for SPY and WQG ternary diagram, respectively. In both cases, the experimental error was modeled by a priori expectation of:

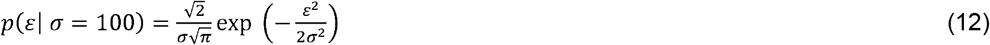

The data *F*(*x*) was then modelled as it were normally distributed with scale parameter ε:

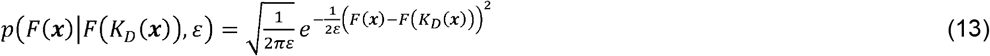

The model was implemented using the pymc (version 5.25) python library ^47^. The model was scaled by the default jitter + adapt_diag method before sampling the posterior parameter space, and a total of 12000 samples were collected in four Markov chain Monte Carlo (MCMC) parallel chains using the No-U-Turn sampling algorithm (NUTS) ^48^. The last 1000 samples from the four chains were combined to yield 4000 samples for the final analysis. For posterior predictions 4000 samples were generated from each of 4000 posterior samples.

## Acknowledgements

This work was supported by grants from the Swedish Research Council (M.I.B., 2017-03025), the Swedish Association Against Rheumatism (M.I.B., R-566961, R-751351, and R-860371), the King Gustaf V:s 80-Year Foundation (M.I.B.), and the regional agreement on medical training and clinical research between the Western Götaland County Council and the University of Gothenburg (M.I.B., ALFGBG-717681 and ALFGBG-965623). S.A.M. acknowledges partial financial support from the Croatian Science Foundation (under the project number HRZZ-IP-2022-10-3456). This project has also received funding from the European Union’s Horizon 2020 research and innovation programme under grant agreement No. 964203 (Long-range electrodynamic INteractions between proteinS — LINkS).

## Author Contributions

G.K., M.I.B. and S.A.M. conceptualized the study outline and discussion and supervised the research. T.N.O., A.L.A., S.A.M., M.I.B. and G.K. analyzed the results from the peptide microarray. M.I.B, J.T., V.Ca., V.Ch., M.P., A.L.A., M.-J. G.-B., J.P., D.M. and G.K. participated in discussions. V.Ch. and M.I.B. tested the web application. The manuscript was prepared by T.N.O., S.A.M. and G.K. with additional input from all authors. All authors reviewed the manuscript.

## Code and data availability statement

The code and microarray data will be deposited in the Zenodo repository upon acceptance of the manuscript. The human proteome clustering web application is openly accessible at http://comproteome.katonalab.eu.

