## Supplemental Online Figures for "Functional Group Composition: The Blueprint for Protein Interactions"

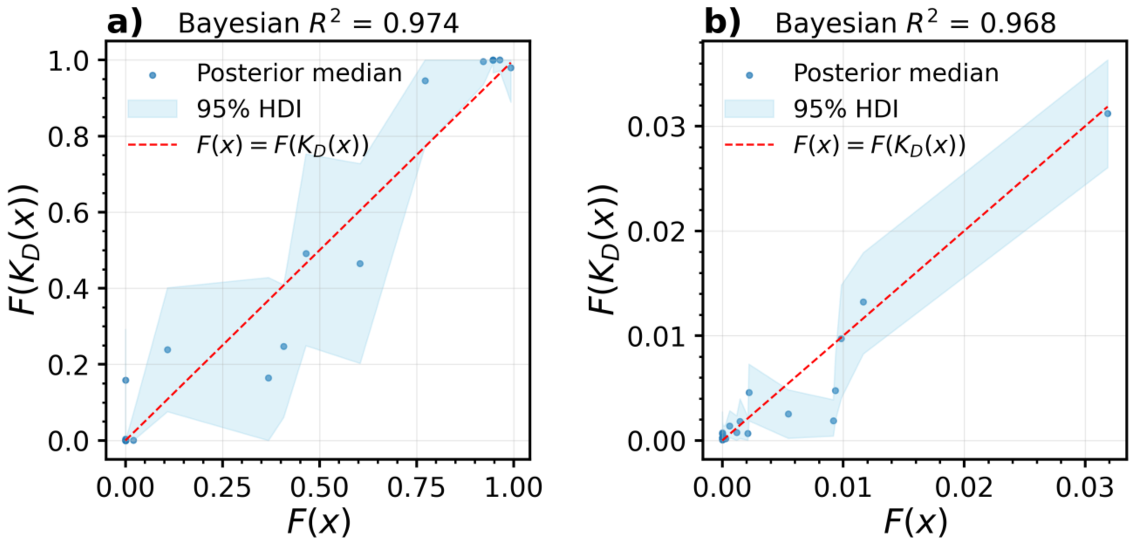


**Figure S1:** Posterior predictive check of the model applied to the a) SPY and b) WQG experiment.

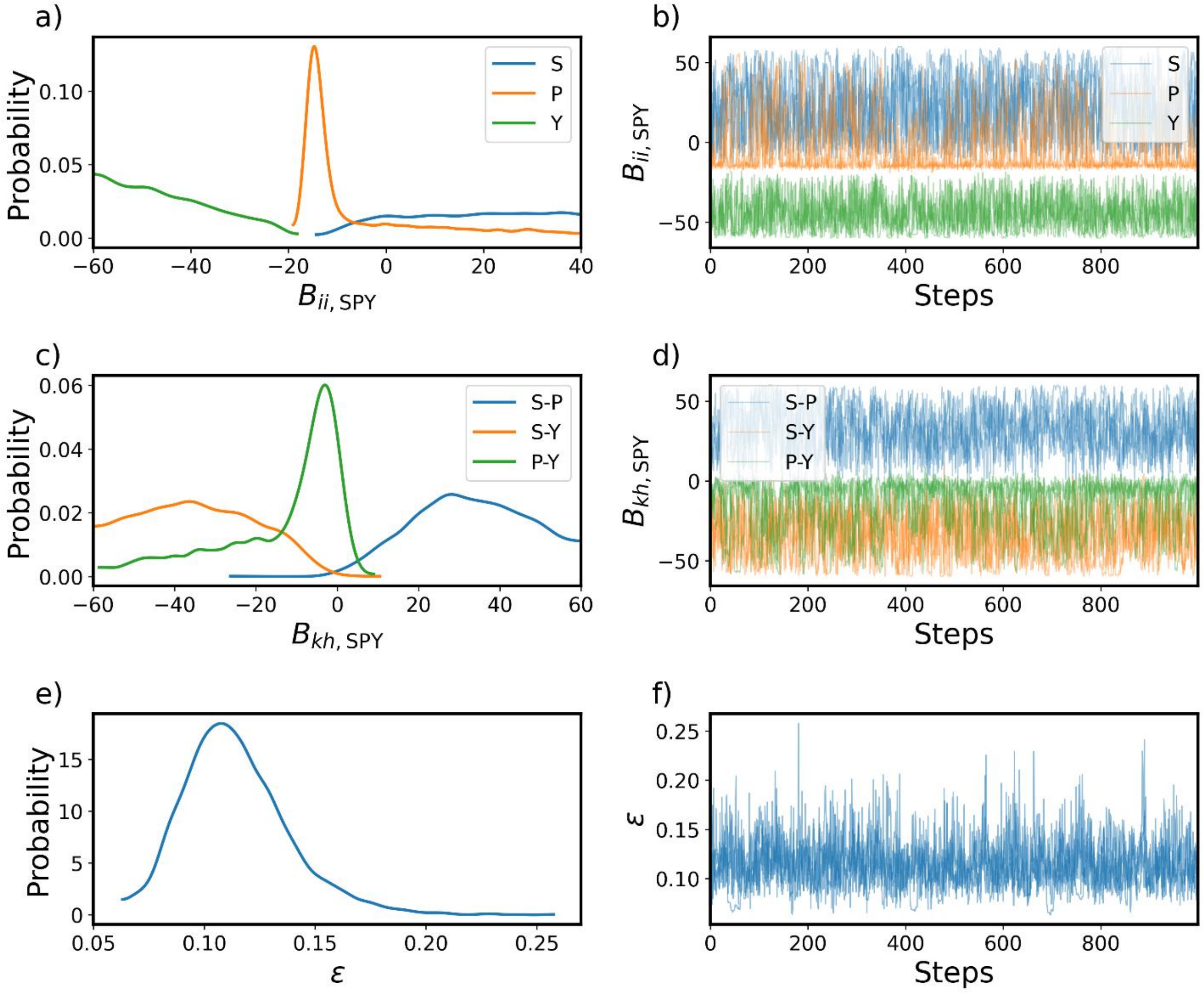


**Figure S2:** Posterior distributions and Markov chain Monte Carlo (MCMC) traces of parameters from the equilibrium thermodynamic model fitted to the SPY experiment.

a) Posterior distribution of self-virial coefficients $\mathbf{B_{ii}}$

b) MCMC traces of $\mathbf{B_{ii}}$

c) Posterior distribution of parameter the cross-virial coefficients $\mathbf{B_{kh}}$

d) MCMC traces of $\mathbf{B_{kh}}$

e) Posterior distribution of parameter $\boldsymbol{\varepsilon}$

f) MCMC traces of $\boldsymbol{\varepsilon}$

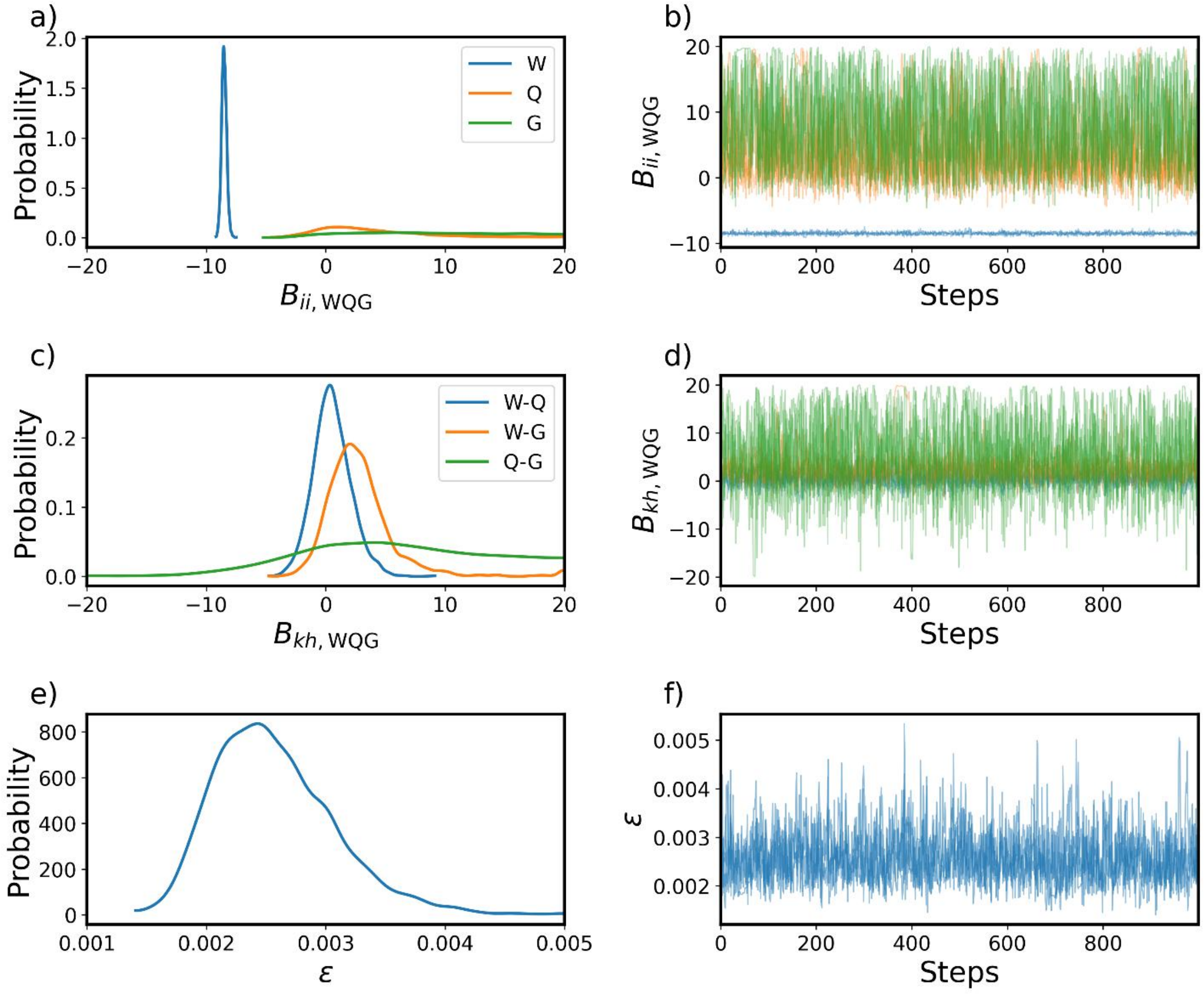


**Figure S3:** Posterior distributions and Monte Carlo (MCMC) traces of parameters from the equilibrium thermodynamic model fitted to the WQG experiment.

a) Posterior distribution of self-virial coefficients $\mathbf{B_{ii}}$

b) MCMC traces of $\mathbf{B_{ii}}$

c) Posterior distribution of parameter the cross-virial coefficients $\mathbf{B_{kh}}$

d) MCMC traces of $\mathbf{B_{kh}}$

e) Posterior distribution of parameter $\boldsymbol{\varepsilon}$

f) MCMC traces of $\boldsymbol{\varepsilon}$

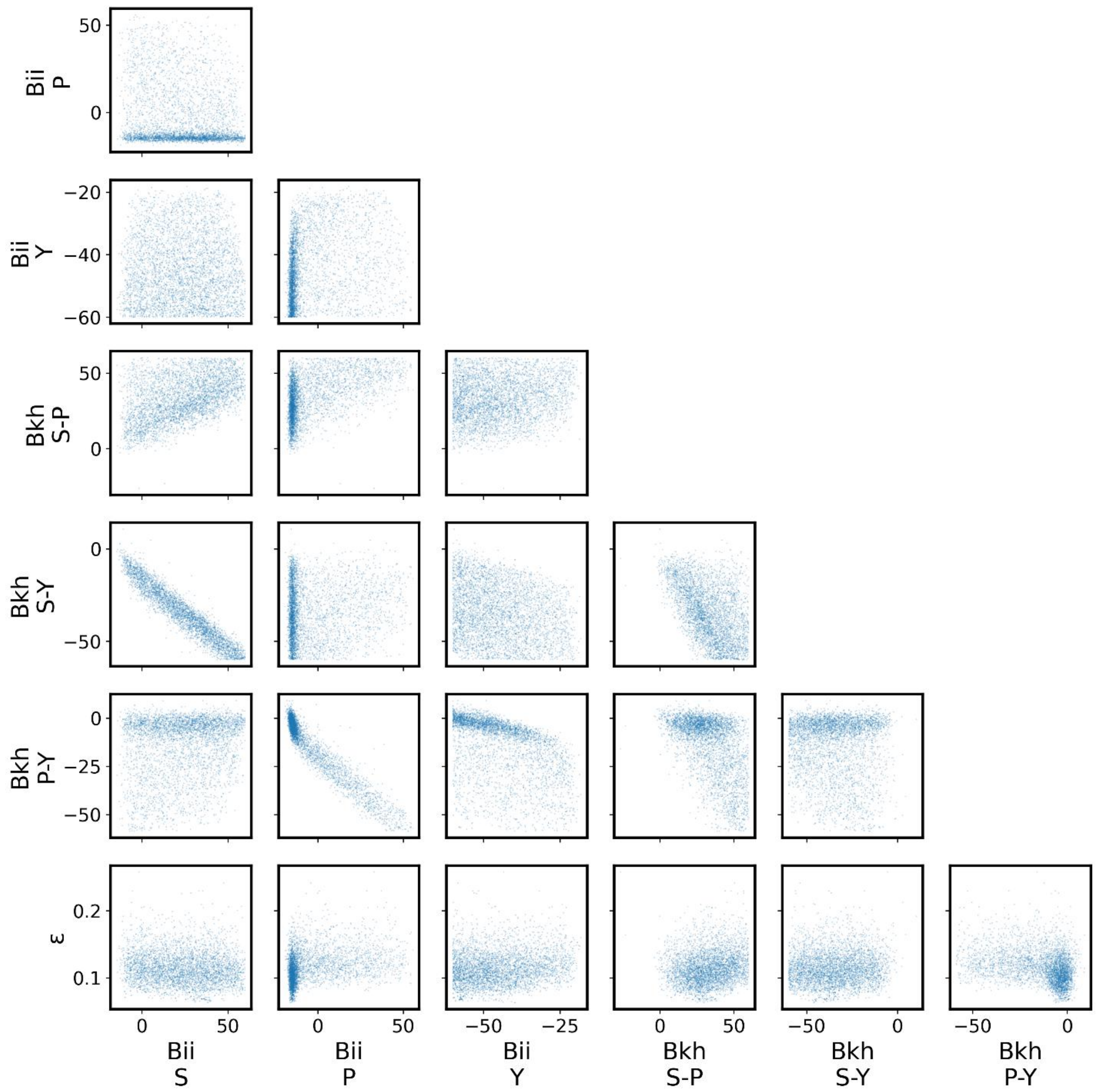


**Figure S4:** Pair plot of parameter traces from the SPY experiment.

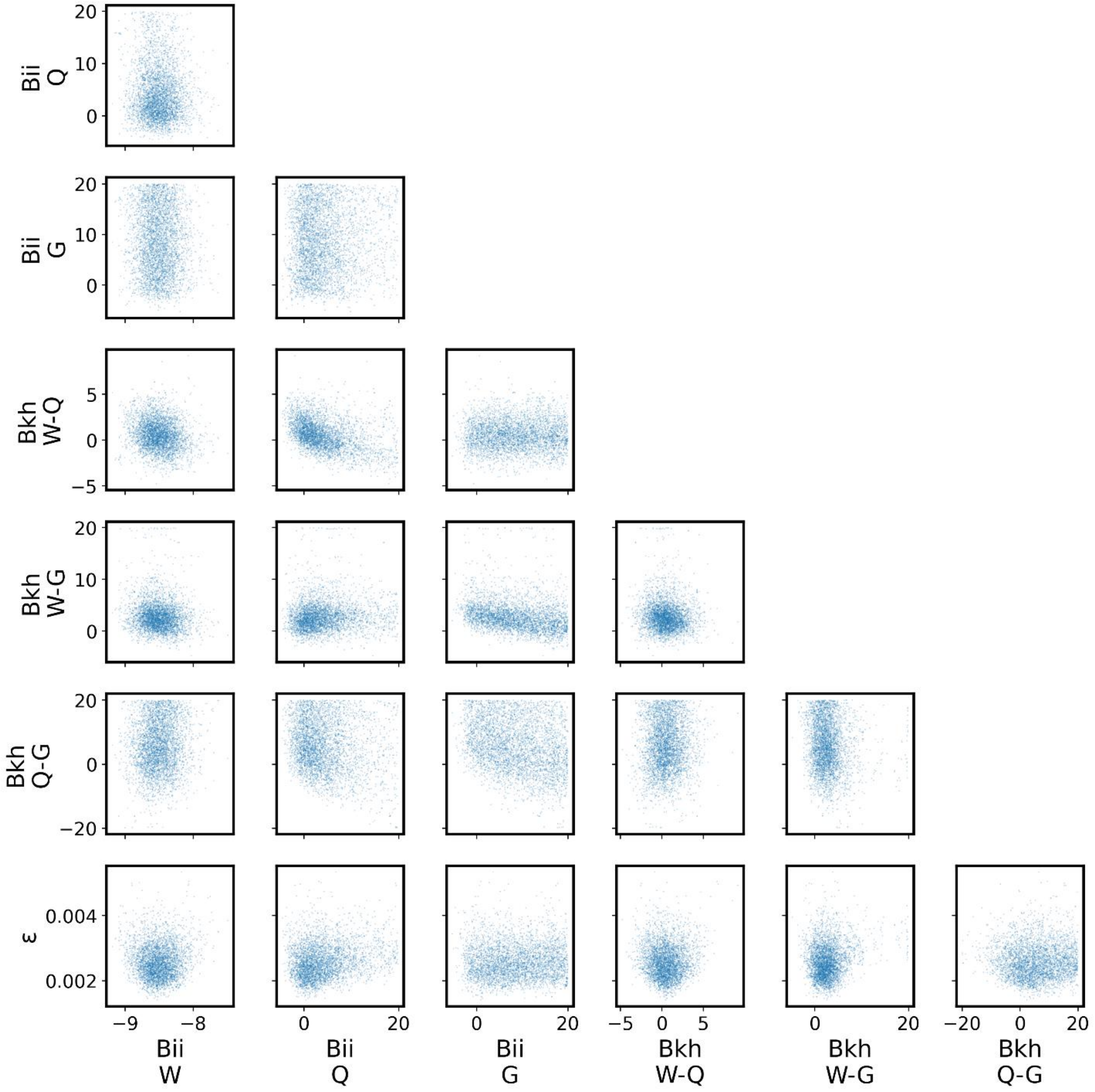


**Figure S5:** Pair plot of parameter traces from the WQG experiment.

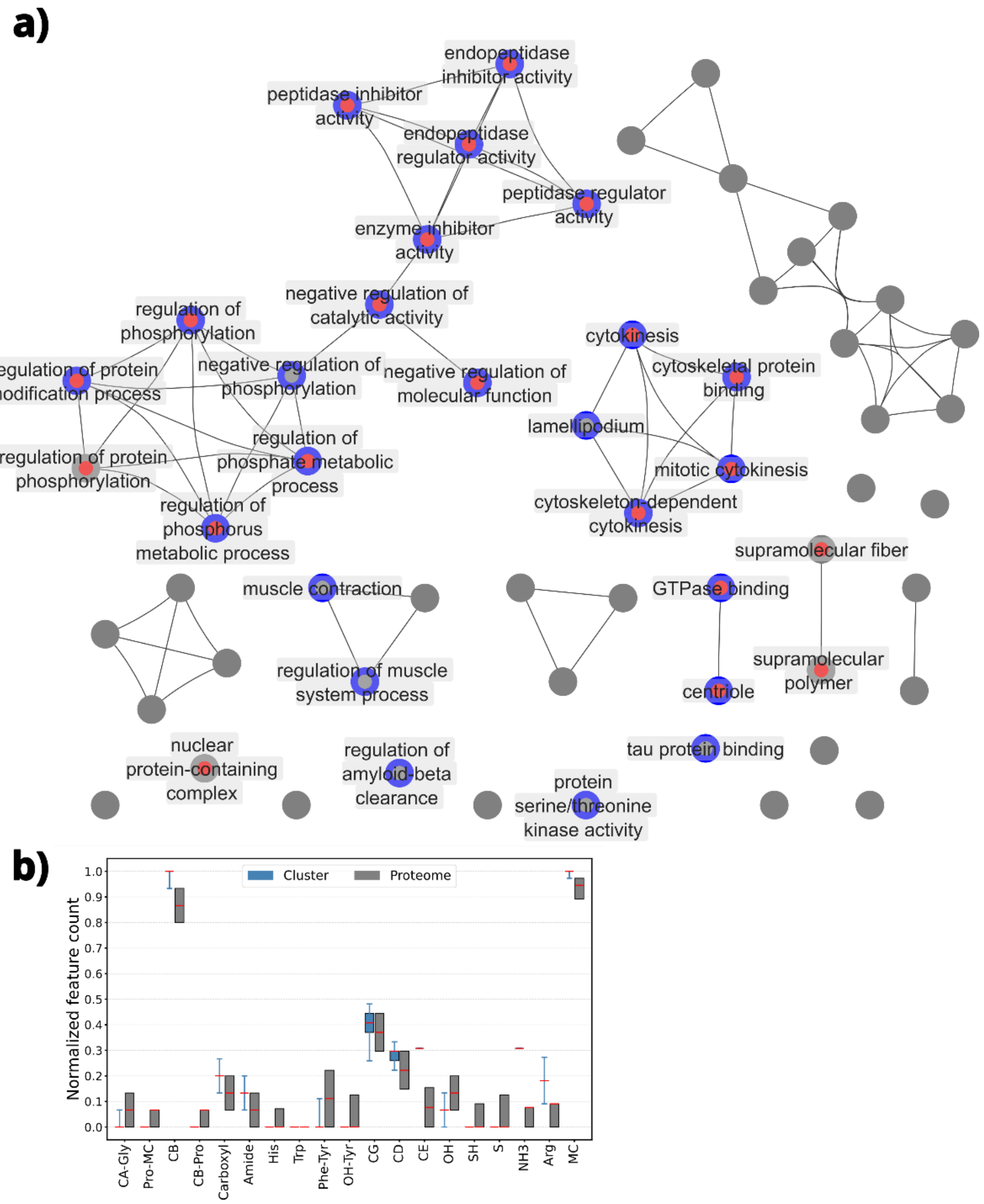


**Figure S6:** Proteins enriched in a fine-grained cluster (k=200,000) containing the $BIRC5_{106\text{-}120}$ region. The cluster contains 37 protein segments (94.6% predicted to bind to survivin) belonging to 35 proteins. a) Functional enrichment network of proteins. Red and blue indicates survivin and ROCK1, respectively. SGO2 is not part of this fine-grained cluster. The p-value cutoff was set at 0.05, and edges were drawn only for node intersections with a Dice similarity greater than 0.7. GO terms (nodes) associated only with a single protein were excluded. b) Boxplot of the normalized composition of protein segments within the cluster (blue) and within the human proteome (gray). The median is indicated by a red mark; whiskers represent the minimum and maximum of the cluster. The minimum and maximum values for proteome segments are 0 and 1, respectively.

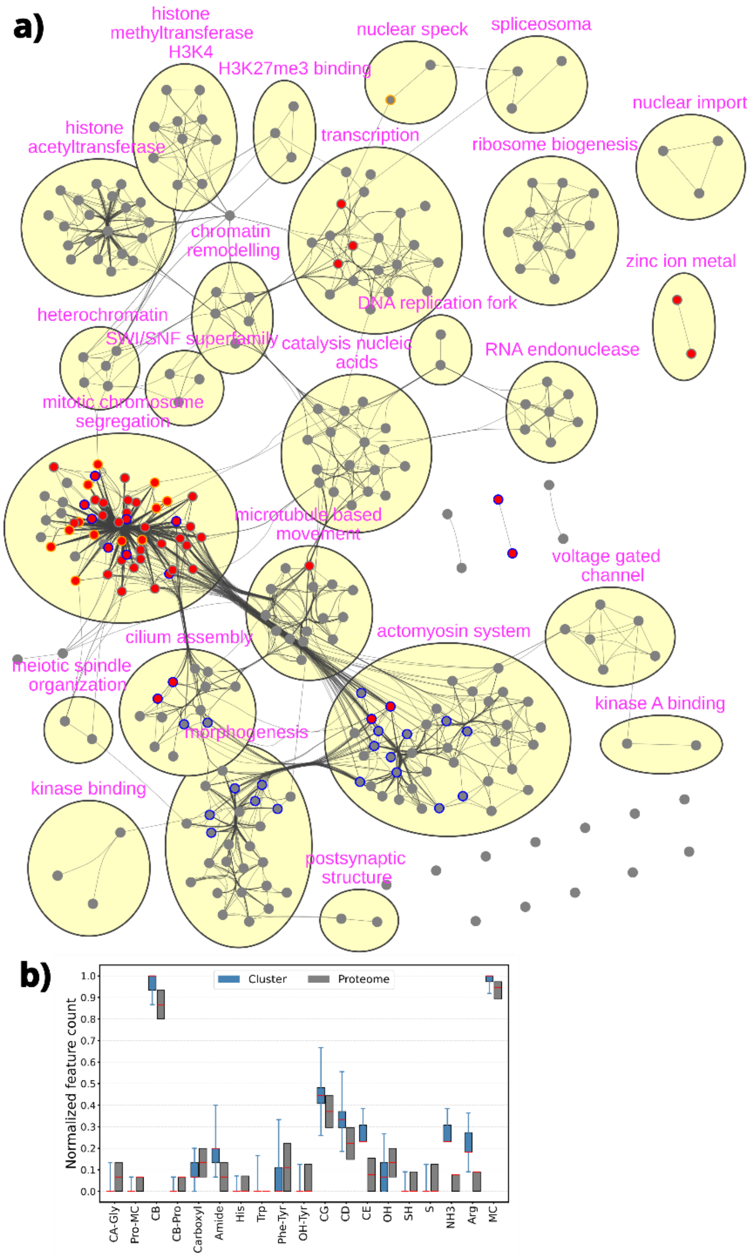


**Figure S7:** Proteins enriched in a coarse-grained cluster (k=2,000) containing the $BIRC5_{106-120}$ region. The cluster contains 1137 protein segments (99.6% predicted to bind to survivin) belonging to 942 proteins. a) Functional enrichment network of proteins. Red, orange and blue indicate survivin, SGO2 and ROCK1, respectively. The p-value cutoff was set at 0.05, and edges were drawn only for node intersections with a Dice similarity greater than 0.3. b) Boxplot of the normalized composition of protein segments within the cluster (blue) and within the human proteome (gray). The median is indicated by a red mark; whiskers represent the minimum and maximum of the cluster. The minimum and maximum values for proteome segments are 0 and 1, respectively.

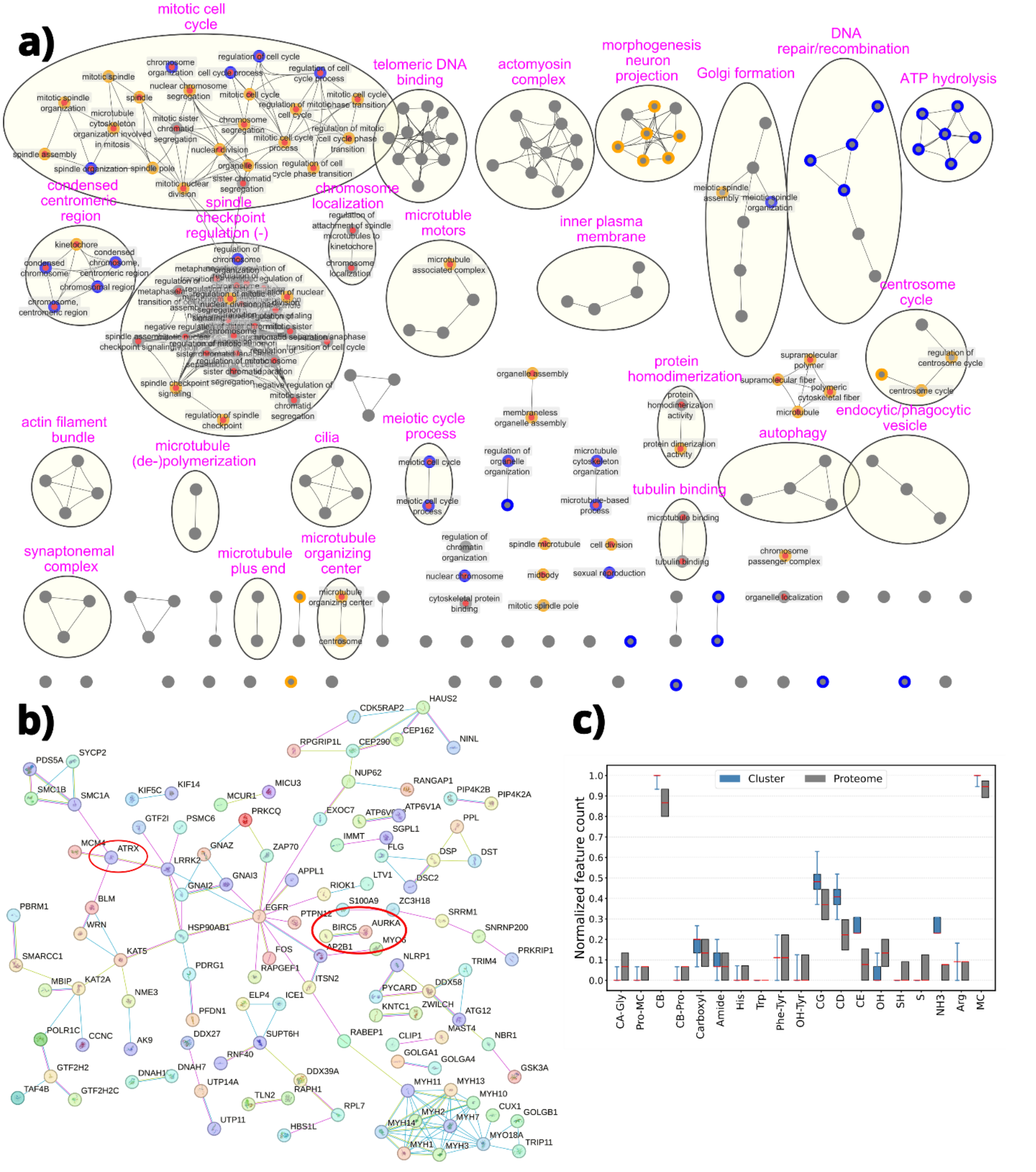


**Figure S8:** a) A functional enrichment of proteins in the cluster that includes BIRC5$_{101-115}$. This includes 304 protein segments (66% predicted to bind to survivin) that belong to 283 proteins. Red, orange and blue indicate survivin, AURKA and ATRX in the intersection with the term, respectively. The p-value cutoff at 0.05, and edges were drawn with similarity >0.7. b) STRING physical subnetwork analysis of the same set of proteins. c) Boxplot of the normalized composition of protein segments within the cluster (blue) and within the human proteome (gray).

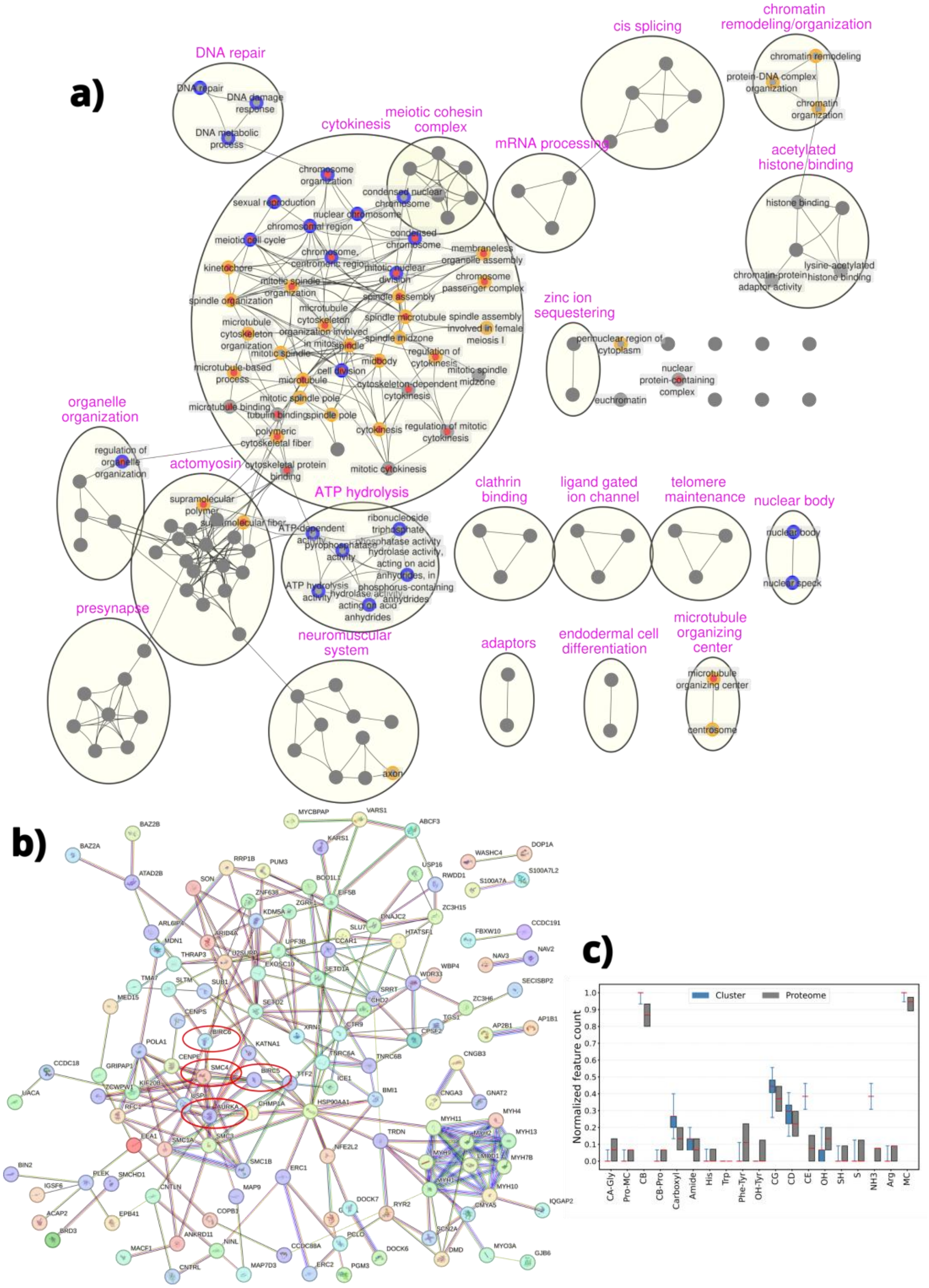


**Figure S9:** a) A functional enrichment of proteins in the cluster that includes both BIRC5$_{111-125}$ and BIRC5$_{116-130}$. This includes 205 protein segments (48% predicted to bind to survivin) that belong to 181 proteins. Red, orange and blue indicate survivin, AURKA and SMC4 in the intersection with the term, respectively. The p-value cutoff at 0.05, and edges were drawn with similarity >0.5. b) STRING physical subnetwork analysis of the same set of proteins. c) Boxplot of the normalized composition of protein segments within the cluster (blue) and within the human proteome (gray).
